# A protein-dependent riboswitch activates ribosomal frameshifting in cardioviruses

**DOI:** 10.1101/2025.07.17.665365

**Authors:** Jemma K. Betts, Cy M. Jeffries, Tim C. Passchier, Hermia C. Y. Kung, Sarah P. Graham, Mahmoud A.S. Abdelhamid, Jamieson A. L. Howard, Timothy D. Craggs, Stephen C. Graham, Ian Brierley, Mark C. Leake, Steven D. Quinn, Chris H. Hill

## Abstract

Programmed −1 ribosomal frameshifting (PRF) is a translational control mechanism used by RNA viruses to regulate the relative abundance of proteins encoded in different reading frames. Cardioviruses exhibit the highest known PRF efficiency, with ∼85% of ribosomes shifting into the −1 frame. This unusual event requires an interaction between the viral 2A protein and a stimulatory element in the RNA genome, but the basis for protein-dependence is unclear. To address this, here we investigate structure and dynamics of the PRF signal in Theiler’s murine encephalitis virus (TMEV). By combining X-ray crystallography, small angle X-ray scattering (SAXS) and single-molecule fluorescence resonance energy transfer (smFRET), we show that 2A binding switches the RNA from a stem-loop conformation into a pseudoknot, and we demonstrate that pseudoknot formation is essential for efficient PRF *in vitro* and in cells. Together, these findings illustrate how the cardiovirus PRF element behaves as a protein-dependent riboswitch, defining the molecular mechanism by which frameshifting is conditionally activated.

## Introduction

Programmed −1 ribosomal frameshifting (PRF) is a translational control mechanism that underpins the replication cycle of hundreds of RNA viruses (reviewed in^1–3^), the collective impact of which are enormous. Examples include HIV-1 (*Lentivirus humimdef1,* ∼1.3 million infections per year^4^), SARS-CoV-2 (*Betacoronavirus pandemicum,* ∼760 million cases and ∼6.9 million deaths^5^), Japanese encephalitis virus (JEV; *Orthoflavivirus japonicum,* ∼100,000 infections and ∼25,000 deaths per year^6^) and porcine reproductive and respiratory syndrome virus (PRRSV; *Betaarterivirus europensis*, ∼$664 million annual loss in the USA^7^).

During a −1 frameshift event, ribosomes are diverted one nucleotide backwards into an alternative reading frame at a specific position on the viral RNA, leading to the translational fusion of polypeptides encoded by the upstream ‘0 frame’ and downstream ‘-1 frame’ sequences^8,9^. The frameshift efficiency is usually fixed, producing a ratio of 0 frame to −1 frame proteins that is optimal for efficient virus replication. Frameshifting is therefore crucial for viral fitness, and sequences directing PRF are highly conserved. Moreover, PRF is not widely used as a gene expression strategy by the host, making it an attractive target for antiviral therapeutics^10–12^.

Frameshifting typically requires two elements. First, a heptameric ‘slippery sequence’ of the form X_XXY_YYZ (where X is any nucleotide, Y is A or U, and Z is any nucleotide except G) permits codon-anticodon base pairing in both 0 (X_XXY_YYZ) and −1 (XXX_YYY) frames. Secondly, a structured RNA ‘stimulatory element’ (usually a pseudoknot or stem-loop) located 5–9 nucleotides downstream acts as a mechanical barrier, resisting unwinding by ribosomal helicase proteins that encircle the mRNA entry channel^13,14^. Ribosomes slow down or pause when they encounter the stimulatory element^15–18^, positioning the slippery sequence in the P and A sites, and causing tension to develop in the RNA^19^. Tandem slippage of tRNAs likely occurs during subsequent translocation attempts (reviewed in^20^), characterised by prolonged rotated hybrid and chimeric states, impaired small subunit head swivel, abnormal elongation factor G (EF-G) function and slow release of E site tRNA^21,22^. During this delay, tRNAs may spontaneously fluctuate between reading frames, reaching an equilibrium based on the relative free energies of pairing in the 0 and −1 frames^22,23^.

RNA stimulatory elements present an intriguing paradox, with many — and often conflicting — features linked to their ability to promote frameshifting. Despite being protein-coding sequences, they must be stable enough to impede the ribosome, whilst also unwinding at physiologically achievable forces during translocation (∼13 pN)^24^. Early work suggested that thermodynamic stability of the stimulatory element would correlate with PRF efficiency^25^, but this is an oversimplification. Subsequent structural work on diverse viral PRF pseudoknots revealed complex features hypothesised to be important, including minor groove triplex interactions between stem 1 (S1) and loop 3 (L3)^26–30^, unusual geometry such as base triples, quadruples or bending at the S1/S2 junction^31–33^ and ‘threading’ of the 5′ end in the SARS-CoV-2 pseudoknot^34–36^. Recently, a cryo-EM study revealed specific interactions between the SARS-CoV-2 pseudoknot and components of the 80S ribosomal helicase^37^, providing evidence for the importance of specific RNA structures. Alternatively, it has been proposed that frameshifting is driven by the existence of multiple interconverting conformations that are sampled by a stimulatory element^38–41^. These ideas can be difficult to reconcile with high-resolution structures of stimulatory elements, which are necessarily static snapshots of a single RNA conformer. This is further confounded by the inaccuracy of deep-learning-based RNA structure prediction tools^42^.

Another layer of complexity is presented by cellular and viral proteins, which can either inhibit^43–45^ or stimulate^46–49^ PRF. A striking case of the latter occurs in the cardioviruses (family *Picornaviridae*), with 85% of ribosomes shifting into the −1 frame at a conserved G_GUU_UUU slippery sequence in Theiler’s murine encephalitis virus (TMEV; *Cardiovirus theileri*)^50–52^ (**Figure 1A-C**). This is the highest known efficiency viral PRF event in nature, but is unusual for several reasons. First, it is entirely conditional on the presence of the viral 2A protein, which binds to an RNA stimulatory element located 13–14 nt downstream^50,51^. Secondly, although this element is predicted to form a seven base pair stem-loop, none of the essential nucleotides reside within the stem. Instead, a pair of guanines in the flanking 5′ region and a cytosine triplet in the loop are required for 2A binding and frameshifting^52^. Finally, the positioning of this element deviates significantly from the canonical 5–9 nt spacer — seemingly too far away to pause the ribosome over the slippery sequence. We previously hypothesised that the cardiovirus stimulatory element may instead form an RNA pseudoknot, which 2A selectively recognises^52,53^. However, this does not adequately explain the protein-dependency of PRF, and structures of 2A proteins from TMEV and encephalomyocarditis virus (EMCV; *Cardiovirus rueckerti*) reveal no homology to known RNA-binding folds^52,53^. Thus, the molecular mechanisms underpinning cardiovirus PRF remain elusive.

**Figure 1.**
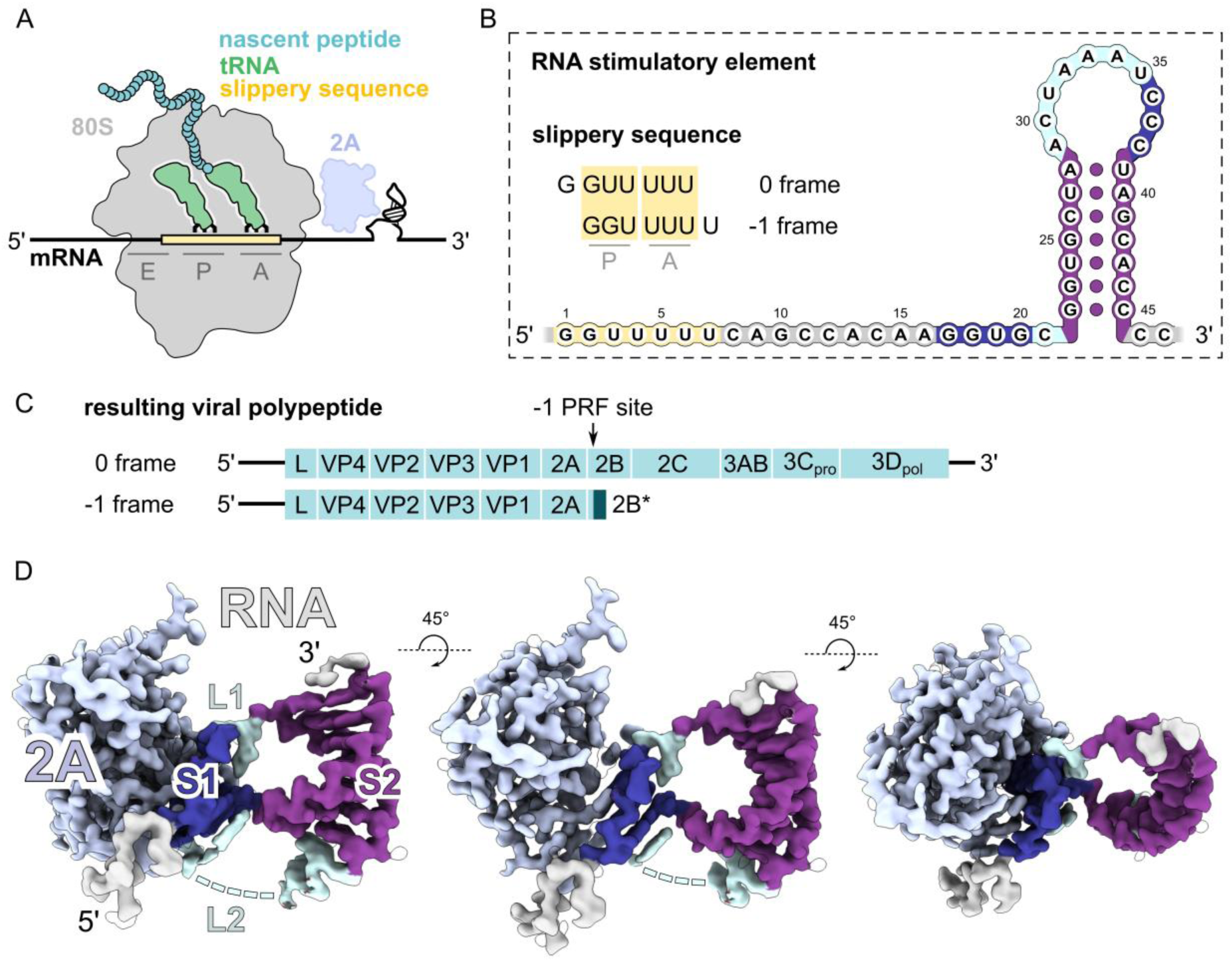
The PRF stimulatory element from TMEV is a protein-pseudoknot complex. A) Diagram of −1 PRF in the cardioviruses. Frameshifting is conditional on the virally encoded 2A protein. The 2A protein (light blue) binds to the structured RNA stimulatory element (black, predicted stem-loop), stalling the elongating ribosome whilst the slippery sequence (yellow) is positioned in the decoding centre. tRNAs (green) may uncouple and recouple in the −1 frame, prior to continuation of the ribosome. B) Details of the TMEV frameshift site. Numbering starts from slippery sequence (yellow). The stimulatory element is predicted to form a 7 bp stem-loop (purple). Conserved GGxG and CCC motifs (dark blue) and divergent interconnecting sequences (cyan) are highlighted. C) The effect of −1 PRF on viral gene expression. Leader protein (L) and capsid proteins (VP4, VP2, VP3, VP1) are encoded 5ʹ to the −1 PRF site (indicated by black arrow) whereas the replicative proteins (2B, 2C, 3AB, 3C_pro_ – protease, 3D_pol_ – RNA polymerase) are encoded 3ʹ to the −1 PRF site. Frameshifting results in the truncation of the viral polyprotein, creating the 2B* product, and downregulating expression of the replicative proteins. Up to 85% of elongating ribosomes are diverted into the −1 frame. D) 1.9 Å X-ray crystal structure of the 2A-RNA complex reveals that the RNA adopts a pseudoknot fold, not a stem-loop, when bound to 2A. Electron density (2F_o_-F_c_) within 1.2 Å of the refined models is shown in three views. 5ʹ and 3ʹ nucleotides not involved in base pairing are shown in grey, stem 1 (S1) in blue, stem 2 (S2) in purple, and loops (L1, L2) in cyan, corresponding to the colouring of the sequence in B). Three disordered nucleotides in L2 are shown as a dashed line.

Here, we combine X-ray crystallography, SAXS and single-molecule fluorescence resonance energy transfer (smFRET) to examine the structure and dynamics of the TMEV stimulatory element in the presence and absence of 2A protein. We first determine a 1.9 Å structure of the 2A-RNA complex, revealing a compact RNA pseudoknot with a non-canonical two-stem architecture. We next demonstrate that the RNA stimulatory element behaves as a protein-dependent riboswitch, undergoing a dramatic conformational rearrangement from a seven base pair stem-loop into a pseudoknot upon 2A binding. Our structure explains the protein dependence of pseudoknot formation, and we dissect the sequence specificity of this unique molecular switch, *in vitro* and in cells. Together, this work defines the conformational repertoire of a protein-dependent riboswitch and demonstrates that — even though cardiovirus PRF elements are indeed conformationally plastic — one specific structure is likely the major determinant of PRF. We suggest that frameshifting signals in other viruses may be similarly regulated by as-yet-unidentified proteins or ligands, explaining some of the reported conformational heterogeneity.

## Results

### The PRF stimulatory element from TMEV is a protein-pseudoknot complex

To understand the molecular basis of PRF in the cardioviruses, we began with structural studies of the stimulatory element from TMEV. We determined the X-ray crystal structure of a complex between TMEV 2A protein (amino acids 1–133, corresponding to 923–1055 of the TMEV polyprotein, UniProt: <u>P08545</u>) and a minimal RNA stimulatory element (nucleotides 14– 47 in **Figure 1B**, corresponding to 4257–4290 of the TMEV genome, GenBank: <u>NC_001366.1</u>). The structure was refined at 1.9 Å resolution (PDB: <u>9RVP</u>, **Table 1**) revealing that the RNA adopts a pseudoknot conformation when bound to 2A protein (**Figure 1D** and **<u>movie S1</u>**). 2A adopts the same globular ‘beta-shell’ architecture as the *apo* structure^53^ (**Figure S1A**, C_α_ backbone RMSD of 0.39 Å over 125 residues).

**Table 1.** Crystallographic data collection and refinement statistics^a^.

|  | 2A-RNA pseudoknot<br>PDB ID: 9RVP |
| --- | --- |
| Data collection |  |
| Wavelength (Å) | 0.9537 |
| Space group | C 121 |
| Cell dimensions |  |
| *a*, *b*, *c* (Å) | 104.19, 45.93, 65.72 |
| $\alpha$, $\beta$, $\gamma$ (°) | 90.00, 127.34, 90.00 |
| Resolution (Å) | 52.25-1.90 (1.94-1.90) |
| *R*<sub>merge</sub> | 0.152 (0.728) |
| *I* / σ*I* | 7.1 (3.3) |
| Completeness (%) | 98.7 (98.3) |
| Multiplicity | 6.5 (6.7) |
| CC<sub>1/2</sub> | 0.974 (0.640) |
| Refinement |  |
| Resolution (Å) | 52.25-1.90 (2.02-1.90) |
| No. reflections | 19410 (3175) |
| *R*<sub>work</sub> / *R*<sub>free</sub> | 0.1931 / 0.2295 |
| No. atoms |  |
| Protein<sup>b</sup> | 2043 |
| RNA<sup>b</sup> | 962 |
| Water | 249 |
| *B*-factors |  |
| Protein | 28.1 |
| RNA | 47.7 |
| Water | 37.2 |
| R.m.s. deviations |  |
| Bond lengths (Å) | 0.009 |
| Bond angles (°) | 0.831 |
| Ramachandran statistics |  |
| Preferred (%) | 98.35 |
| Allowed (%) | 1.65 |
| Outliers (%) | 0.00 |
| All-atom clashscore | 1.33 |
| Rotamer Outliers (%) | 0.00 |

<sup>a</sup>Data were recorded from a single native crystal. Values in parentheses are for highest-resolution shell. <sup>b</sup>Including hydrogen atoms.

The RNA pseudoknot comprises two non-coaxially stacked stems in a side-by-side arrangement, with a tilt of ∼55° between their helical axes. (**Figure 2** and **Figure S1B**). Stem 1 (S1) consists of three Watson-Crick G-C base pairs between GGxG and CCC triplet motifs, providing a structural basis to the previous biochemical characterisation^49–53^. Interestingly, the three guanosine nucleotides (G17, G18 and G20) comprising S1 are not consecutive, and U19 is flipped out by S1 formation. This results in an unusual helical geometry: the first two G-C base pairs (G17-C38 and G18-C37) exhibit a typical helical twist of ∼36.2°, but the final base pair (G20-C36) has a twist of ∼50.6° relative to G18-C37 (**Figure S1C**), causing compression of the major groove.

**Figure 2.**
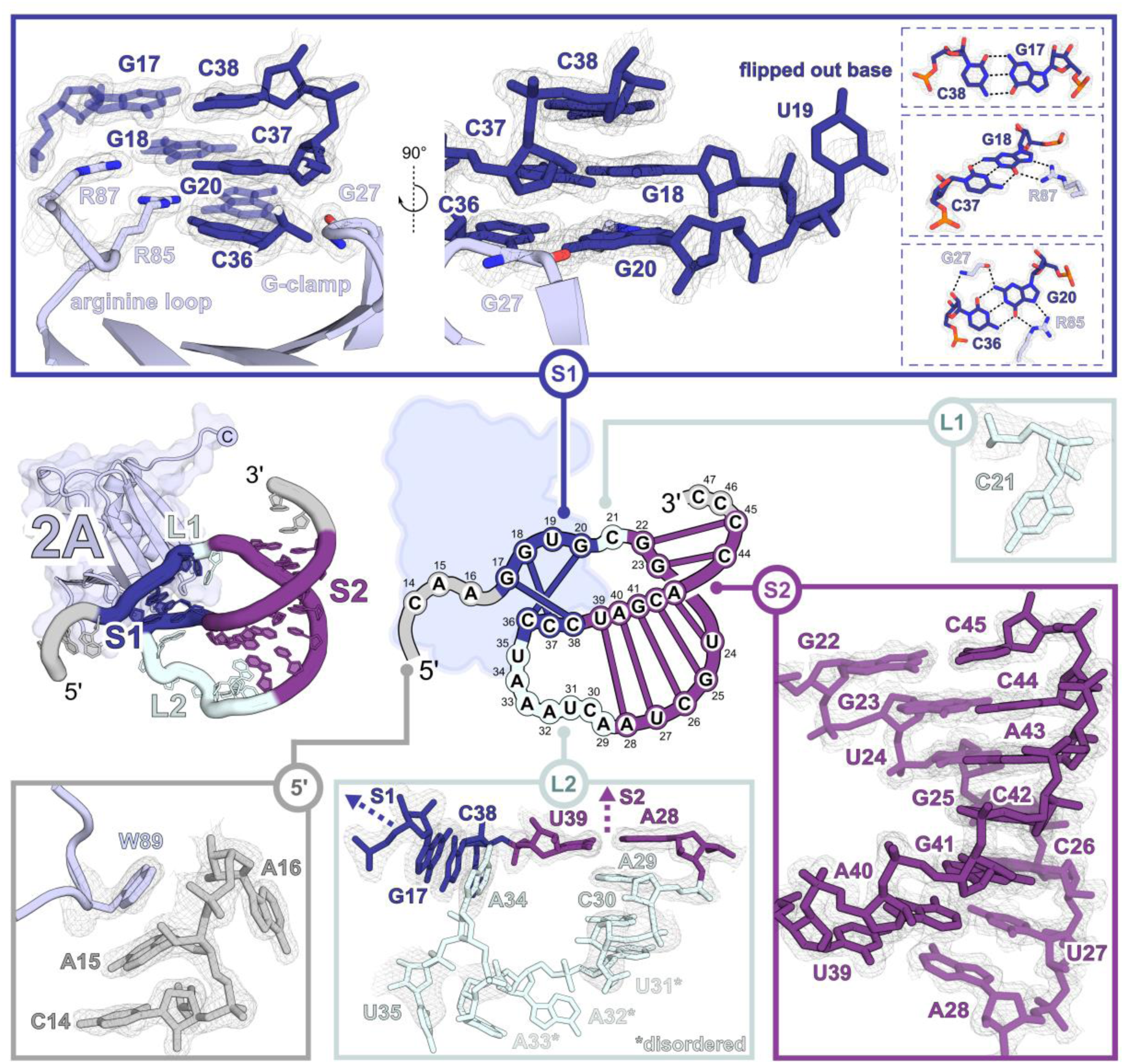
Structural features of the RNA pseudoknot in complex with 2A. Close-up views illustrating key regions of the RNA pseudoknot (also see **<u>movie S1</u>**). The structure is shown as sticks, modelled into 2F_o_-F_c_ electron density (grey mesh) contoured at 0.64 e^-^/Å^3^ (2.0 RMSD). (*Centre*) Topological diagram of the pseudoknot with base identity indicated. S1 consists of three G-C Watson-Crick base pairs comprised of the conserved GGxG and CCC triplet motifs, supported by the arginine loop (R85 and R87) and ‘G-clamp’ (G27) of 2A. The flipped base (U19) allows the formation of S1. Details of hydrogen bonding in S1 are shown. L1 is a single nucleotide (C21). S2 is a seven Watson-Crick base paired A-form RNA helix. L2 is mostly disordered. The 5ʹ end of the RNA stacks against W89.

Such geometry is unfavourable, but S1 is stabilised by the insertion of the 2A ‘arginine loop’ into the major groove (**Figure 2** and **<u>movie S1</u>**). The guanidinium groups of R87 and R85 are approximately coplanar with the second (G18-C37) and third (G20-C36) base pairs of S1 respectively, forming two ‘arginine forks’ — recurrent structural motifs in protein-RNA interactions^54–56^. R87 stabilises the second base pair of S1 (G18-C37) via interactions between Nη and the Hoogsteen edge of G18, and a hydrogen bond between Nε and a ribonucleotide phosphate oxygen of G17 (HηηP fork). R85 stabilises the third base pair of S1 (G20-C36) via interactions between Nη2 and Nε atoms and the Hoogsteen edge of G20, and between Nη2 and the ribose 2ʹ hydroxyl of G18 (HS fork). On the other side of G20, the carbonyl oxygen of glycine 27 forms a hydrogen bond with N2, such that this guanine base is “clamped” between R85 and glycine 27. Comparison to the *apo* structure of TMEV 2A illustrates how glycine 27 moves 1.6 Å towards G20 while the guanidinium group of R85 rotates: we therefore refer to this as the “G-clamp” interaction (**Figure S1A**). This third base pair (G20-C36) additionally packs against the 2A beta-sheet, comprising R7, D9, F11 from β1, H67 from β4 and K24 from β2, the latter of which forms a hydrogen bond with the ribose 2ʹ hydroxyl of G20. Immediately prior to S1, the 5′ end of the stimulatory RNA also interacts with the surface of 2A, with A15 stacking against W89 (**Figure 2**). Together, these interactions bury a surface of 634 Å^2^. AlphaFold3^57^ was unable to predict this interface, instead suggesting interactions between 2A protein and S2 (**Figure S1D**).

Downstream of S1, loop 1 (L1) is a single nucleotide (C21) leading into stem 2 (S2), a 7 bp A-form helix (C22–A28; U39–G45) (**Figure 2**), consistent with previous bioinformatic predictions and structure probing experiments^50,51^. S2 is connected to the 3′ strand of S1 by loop 2 (L2, A29–U35, **Figure 2**). However, the electron density for L2 is poor (**Figure 2**) demonstrating that U31, A32 and A33 are flexible or disordered. Nevertheless, A29 and C30 stack against A28 from S2, and weak density suggests that A34 may stack against C38 from S1. Loop 3, a feature of some H-type pseudoknots^58^, is entirely absent.

### The RNA stimulatory element switches conformation upon 2A binding

We next examined to what extent our crystal structure reflects the conformation of 2A protein and RNA in solution. To do this, we analysed 2A protein, RNA, and the 2A-RNA complex separately by size exclusion chromatography coupled to small angle X-ray scattering (SEC-SAXS, **Figure S2A-F, Figure 3A-B** and **Table 2**). All samples were monodisperse, allowing determination of the radius of gyration (*R*_g_), maximum particle dimension (D_max_), and the calculation of multiple molecular envelopes and atomistic models describing the data (**Figure S2** and **Table 2**). The 2A protein sample (**Figure S2A**, *R*_g_ = 1.89±0.05 nm) matched the protein component of the crystal structure, with an ensemble of five models (**Figure S2B**, χ^2^ = 0.92– 0.97) illustrating the conformational flexibility of disordered N- and C-termini. Similarly, the 2A-RNA complex sample (**Figure 3A** and **Figure S2E**, *R*_g_ = 2.23±0.02 nm) was in good agreement with the crystal structure (**Figure S2F**, five models, χ^2^ = 1.11–1.35), confirming that the pseudoknot-protein complex likely forms in solution. However, the scattering data from the RNA-only sample (**Figure S2C**, *R*_g_ = 1.91±0.02 nm) could not be explained by the pseudoknot component of the crystal structure (**Figure S2G**, χ^2^ = 13.9) and attempts to further refine the RNA pseudoknot to match the SAXS data were unsuccessful. Instead, these data are explained by a 7 bp stem-loop with a flexible unpaired 5ʹ tail (**Figure 3B** and **Figure S2D**, nine models, χ^2^ = 1.11–1.20), suggesting that the RNA may undergo a large conformational rearrangement upon 2A binding.

**Figure 3.**
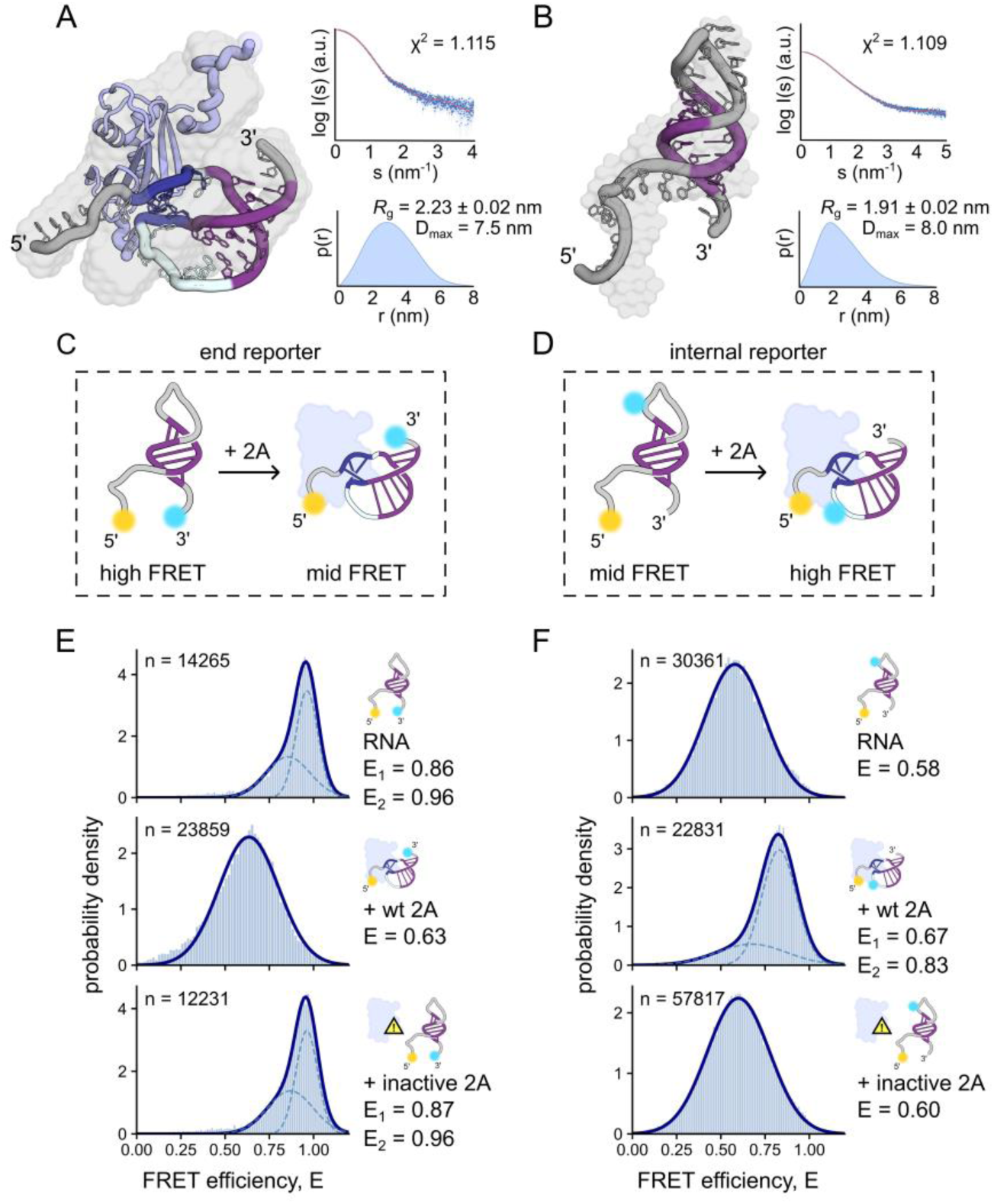
The RNA stimulatory element switches conformation from stem-loop to pseudoknot upon 2A binding. A,B) SEC-SAXS envelope with plots showing alignment between experimentally collected SAXS data (blue circles) and calculated data from structural models (red lines), and the pair-distance distribution P(r) allowing the determination of *R*_g_ ± SD and D_max_. A) 2A-RNA complex with X-ray crystal structure fitted to the data. B) RNA in isolation, supporting a stem-loop model. C,D) Diagrams showing smFRET RNA reporter construct design using the Cy3 (cyan) and Cy5 (yellow) FRET pair. E,F) Corrected FRET efficiency (E) histograms combined from three biological repeats of the E) end reporter and F) internal reporter wild-type RNAs. Showing data for (*upper*) RNA in isolation, (*middle*) RNA in the presence of 0.5 µM wild-type 2A and (*lower*) RNA with 0.5 µM inactive 2A_R85A/R87A_. Corrected average FRET efficiencies (E) and number of molecules (n) are reported in each histogram. Gaussian fits are shown as dashed curves for two Gaussians and a solid blue line for a single Gaussian. R^2^ = 0.99 for all gaussian fits.

**Table 2.**
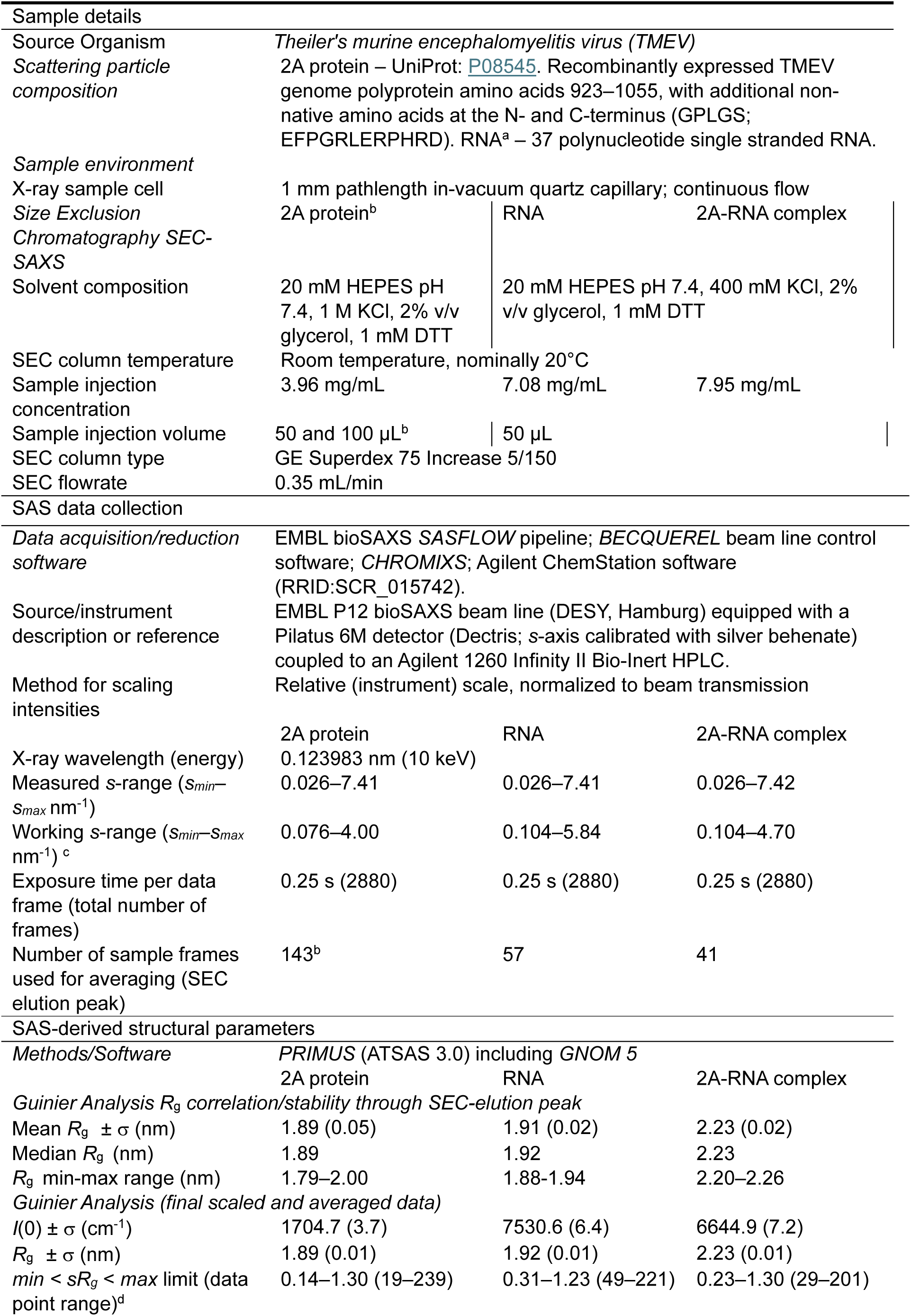
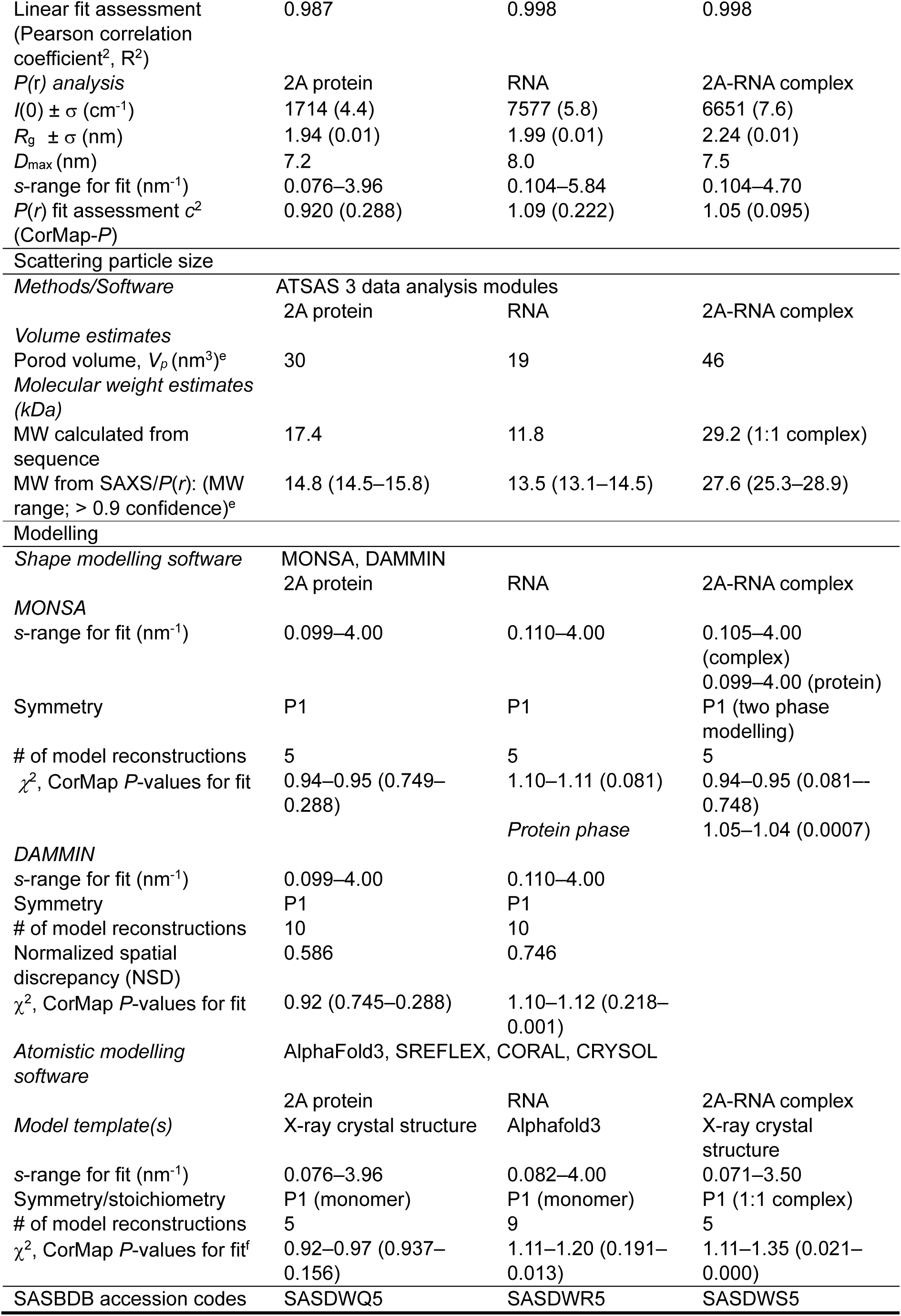
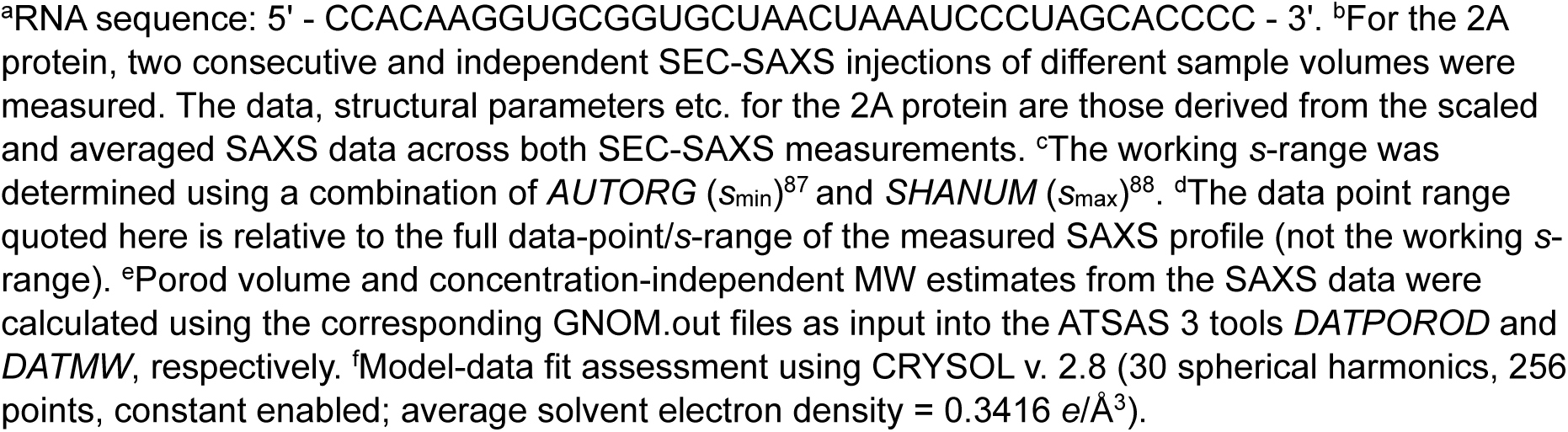
SAXS data collection and analysis parameters: 2A protein, RNA and 2A-RNA complex.

To investigate this further, we conducted smFRET experiments on freely diffusing molecules to explore the conformations sampled by the frameshifting RNA. We designed two FRET reporter RNAs with the Cy5/Cy3 FRET pair: an ‘end reporter’ labelled with 5ʹ Cy5 and 3ʹ Cy3 fluorophores (**Figure 3C** and **Figure S2H**), and an ‘internal reporter’ labelled with 5ʹ Cy5 and an internal Cy3 fluorophore attached to U31 within L2 of the pseudoknot (**Figure 3D** and **Figure S2I**). These were designed to minimally impact RNA folding, avoid dye quenching by adjacent guanine bases, and generate distinct FRET efficiencies in both stem-loop and pseudoknot conformations. The use of alternating-laser excitation (ALEX) confocal imaging^59^ allowed for discrimination between singly- and doubly-labelled FRET-active forms, enabling us to exclusively conduct our downstream analyses on *bona fide* FRET events.

The end reporter RNA exhibited one major high-FRET species (**Figure 3E**, E_2_ = 0.96±0.01, ∼31 Å interdye distance), consistent with a stem-loop conformation. A second minor FRET population (E_1_ = 0.86±0.03, ∼39 Å interdye distance) could indicate mobility of the 5′ single-stranded region in some molecules, consistent with the SAXS data for the RNA alone (**Figures S2C, S2D**). In the presence of 2A, we observe a single mid-FRET population (E = 0.63±0.01) accompanied by a broadening of the FRET distribution (**Table S3**), consistent with the ends moving further apart (∼49 Å) and becoming more mobile. This agrees with our crystal structure, in which neither the 5′ nor 3′ ends of the RNA are stabilised by direct contacts with protein. We do not observe this mid-FRET population in the RNA-only measurements, suggesting that the RNA does not sample the pseudoknot conformation in the absence of 2A.

FRET measurements of the internal reporter RNA revealed a single, mid-FRET population (E = 0.58±0.01) (**Figure 3F**), consistent with a stem-loop conformation (∼50 Å interdye distance). Upon addition of 2A, we observe a shift to a major high-FRET population (E = 0.83±0.01) as the dyes move closer together (∼41 Å) during pseudoknot formation. The FRET distribution also narrows, implying that complex formation stabilises the position of the internal dye. A minor second FRET population is also induced by 2A binding (E = 0.67±0.02, ∼47 Å). To demonstrate that these changes in FRET efficiency are due to specific interactions with 2A protein and not artefacts of molecular crowding, we performed control experiments with inactive 2A_R85A/R87A_ protein harbouring arginine loop mutations. For both reporters, addition of 2A_R85A/R87A_ results in FRET populations identical to RNA-only experiments (**Figures 3E, 3F**). We also confirmed that ionic strength or identity (Na^+^, K^+^, Mg^2+^) have no effect on the conformation of the RNA (**Figure S3**): in the absence of 2A protein, the stem-loop conformation persists, irrespective of salt concentration.

Given that conformational plasticity of stimulatory elements has been proposed to regulate frameshifting^38–41^, we implemented burst variance analysis (BVA)^60^ to further examine dynamics. Briefly, BVA compares the standard deviation of FRET from individual molecules over the timescale of diffusion to the theoretical shot-noise distribution. Our results with both reporters confirm that the RNA does not spontaneously sample multiple stable conformations (**Figure S4**). The end reporter RNA is generally more dynamic than the internal reporter RNA, as indicated by the standard deviation being greater than expected from shot-noise. However, this is likely a feature of dye placement rather than a physiologically relevant characteristic. Overall, our SAXS and FRET analyses support a model in which the stimulatory RNA behaves like a binary switch between two major states: It exists as a stem-loop in isolation, but flips into a pseudoknot in the presence of 2A.

### Pseudoknot formation depends on stem 1 sequence, and stem 2 length

To dissect the molecular rules that govern pseudoknot formation, we created a series of mutant RNAs and assessed their binding to 2A by electrophoretic mobility shift assay (EMSA) and microscale thermophoresis (MST) (**Figures 4A, 4B**). We first tested the importance of the specific arginine-guanine interactions observed in the structure (**Figure 2**). Mutations were designed to preserve S1 base pairing and the unusual helical geometry (including the flipped-out base) but remove the possibility of direct arginine-guanine contacts (**Figure 4C**). These included a complete stem swap, and substitution of G-C pairs with A-U and U-A pairs. As assessed by EMSA and MST, none of these mutants could bind to 2A (**Figure 4C**), demonstrating strong specificity for guanine. We also deleted the flipped-out base (ΔU19) allowing S1 to adopt a more regular conformation. However, this did not detectably bind to 2A, highlighting the importance of irregular S1 geometry.

**Figure 4.**
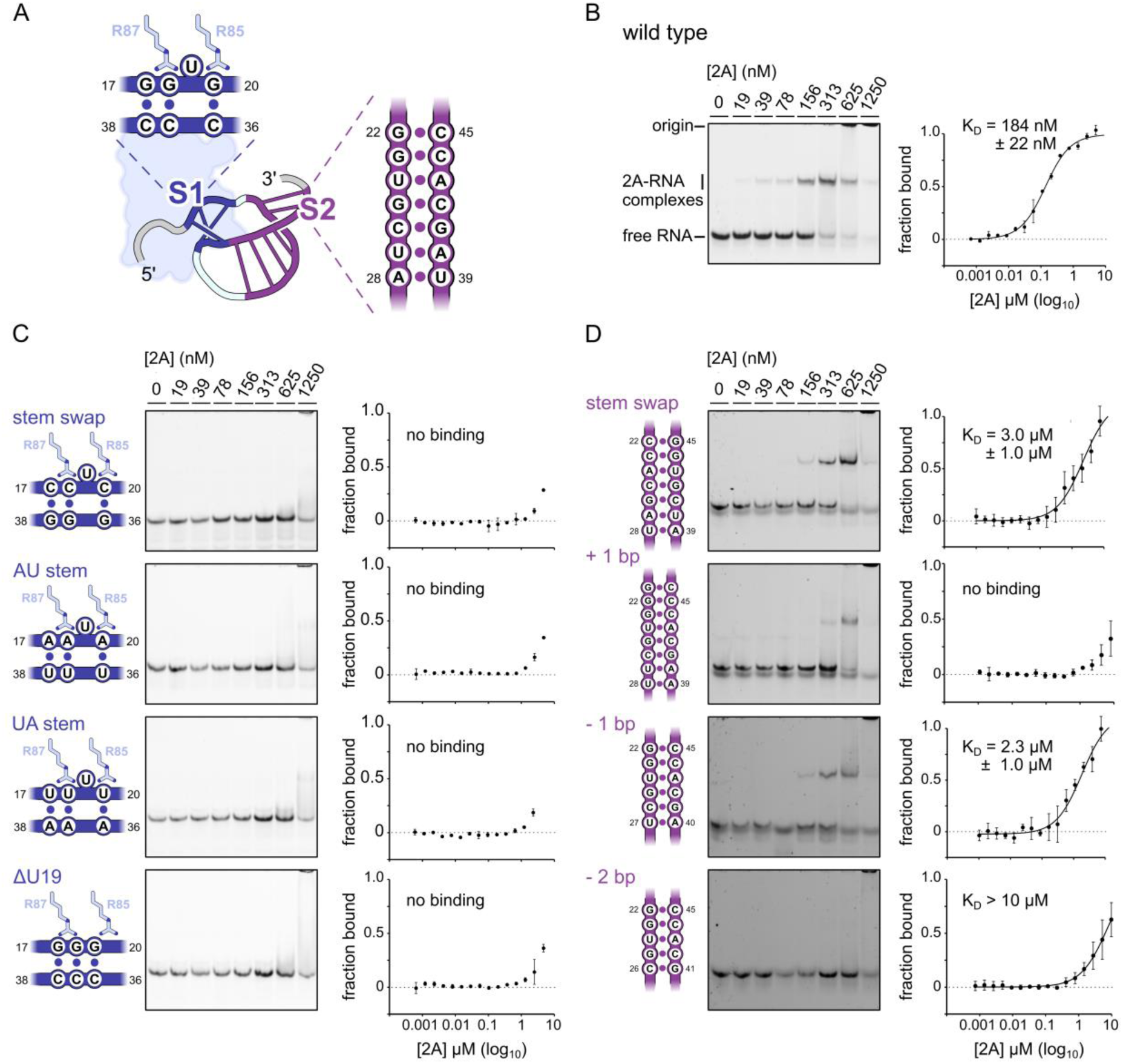
Pseudoknot formation depends on stem 1 sequence, and stem 2 length. A) Diagrams showing the location of mutated nucleotides in stem 1 (S1) and stem 2 (S2). B) (*Left*) EMSA analysis of 2A protein binding to wild-type stimulatory RNA. (*Right*) MST binding curves and approximate K_D_ values using the same RNA, assuming 1:1 binding stoichiometry. Experiments were indepedently performed twice, with error bars indicating ± SD. C) (*Left*) Details of each S1 mutation. (*Middle*) EMSA analysis of the effects of S1 mutations on 2A binding, as in B). (*Right*) MST binding curves and approximate K_D_ values using the same S1 mutant RNAs, as in B). D) (*Left*) Details of each S2 mutation. (*Middle*) EMSA analysis of the effects of S2 mutants on 2A binding, as in B). (*Right*) MST binding curves and approximate K_D_ values using the same S2 mutant RNAs, as in B).

Similarly, we tested the limits of S2 on pseudoknot formation. S2 mutants included a stem-swap, insertion of an additional base pair, and deletion of 1 or 2 base pairs (**Figure 4D**). The stem-swap RNA could bind 2A with similar affinity to wild-type RNA (**Figure 4D**, K_D_ ∼313 nM, assessed by EMSA). Addition or removal of 1 bp from S2 (creating a 6 or 8 bp S2) had modest effects (∼313–625 nM K_D_, assessed by EMSA) but shortening S2 to 5 bp prevented 2A binding, suggesting that length rather than sequence of S2 is important, with 7 bp optimal but 6–8 bp tolerated. The MST data showed a similar trend, but reported lower affinities than EMSA (**Figure 4D**), possibly due to sample aggregation or interactions with capillary surfaces at micromolar concentrations.

Finally, we tested the importance of 2A residues other than R85 and R87 at the pseudoknot interface (**Figure 2** and **Figure S5A**). H26A and R7A mutations had a mild impact on RNA binding (**Figure S5B, S5C**). K24A and W89A exhibited stronger effects, consistent with the specific RNA contacts at these positions, although smFRET experiments showed that each of the four mutations could still trigger pseudoknot formation (**Figure S5D**). Therefore, the only absolute protein requirement is the conserved 2A arginine loop.

### Disruption of stem 1 formation inhibits ribosomal frameshifting *in vitro* and in cells

To confirm the mechanistic relevance of our structural observations, we next tested the effects of these mutations on −1 PRF during translation of reporter mRNAs, where ribosomes will encounter the pseudoknot-2A complex during elongation (**Figure 5**). All mutations to S1 and S2 were made in the context of nucleotides 4137–4307 of the TMEV genome, comprising the stimulatory element, slippery sequence and ∼35 codons upstream to capture any contributions from the nascent peptide. First, we carried out translation *in vitro* using rabbit reticulocyte lysate (**Figures 5A, 5B**). As expected, PRF was 2A-dependent and highly efficient for the wild-type sequence (91.7±2.7%). All S1 mutants reduced PRF efficiency to background levels (<2%) except ΔU19, which retained some activity (13.8±1.7%), albeit still 2A-dependent. In contrast, S2 mutants had minor effects on frameshifting. The stem-swap (89.9±1.7%) and shortening by −1 bp (87.7±5.8%) led to insignificant changes, whereas lengthening by +1 bp (48.9±21.3%) or shortening by −2 bp (36.9±4.0%) caused larger decreases, consistent with trends in 2A-binding affinity (**Figure 4**). We also tested the effects of 2A protein mutations H26A, R7A, K24A and W89A on PRF (**Figure S5E-G**). Only minor effects were observed (59.1–82.1% PRF efficiency), consistent with the ability of these variants to bind to wild-type RNA (**Figure S5B-D**).

**Figure 5.**
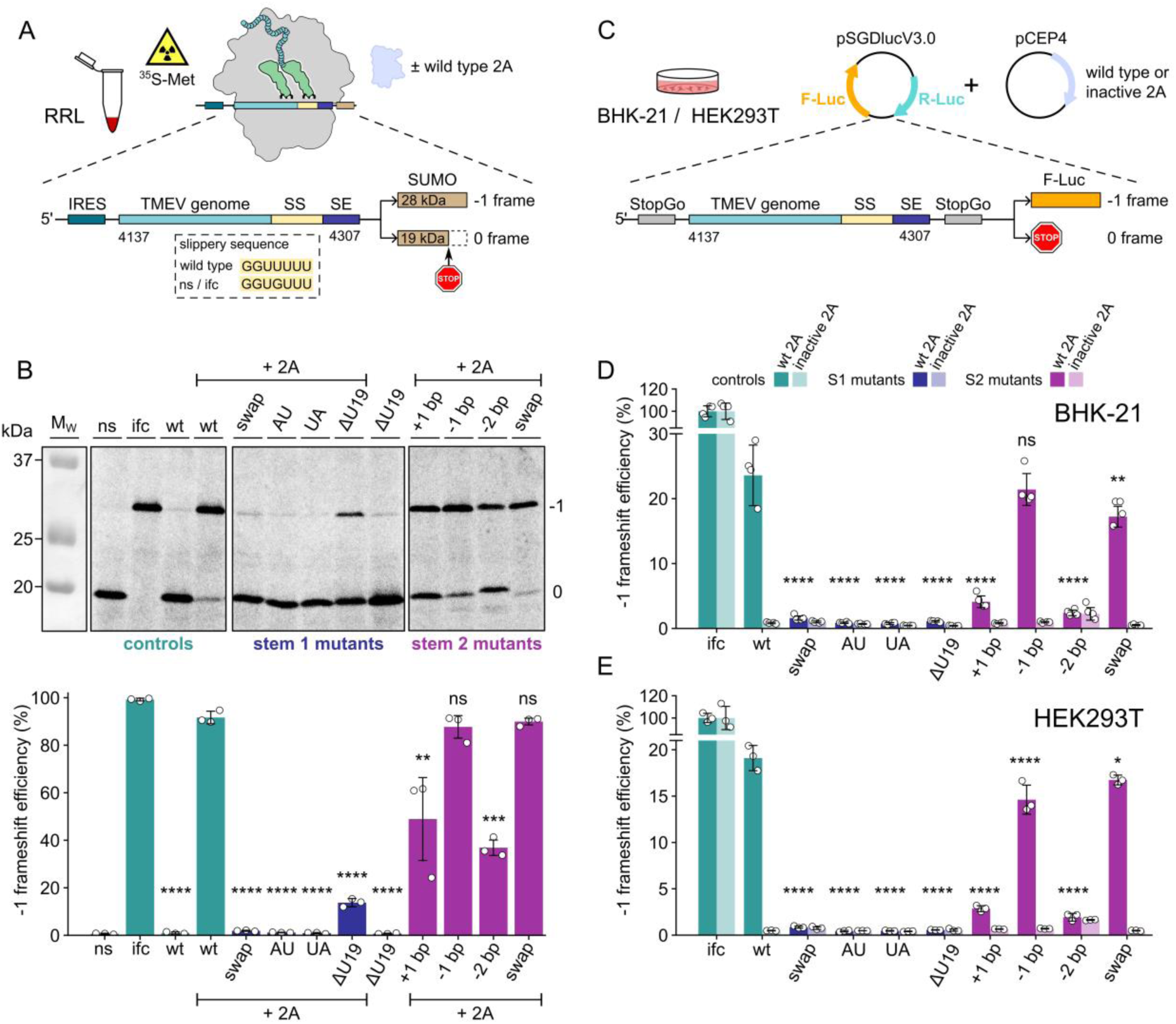
The stem 1 sequence is essential for −1 PRF during translation *in vitro* and in cells. A) Diagram of *in vitro* translation assay to measure frameshifting efficiency. Translation reactions were carried out in rabbit reticulocyte lysate (RRL) in the presence of ^35^S-Met. Reporter mRNAs include 35 codons upstream of the frameshift site (light blue), the slippery sequence (SS, yellow) stimulatory element (SE, dark blue) and SUMO coding sequence (brown) in the −1 frame. B) *(Upper)* SDS-PAGE visualised by autoradiography, showing effects of stem 1 and 2 mutations on frameshifting efficiency *in vitro*. (*Lower*) Densitometric quantification of −1 frameshift efficiencies, corrected for methionine content. Data are represented as the mean ± SD error bars (*n* = 3 biological repeats). ns, not significant, \**p* ≤ 0.05, \*\**p* ≤ 0.01, \*\*\**p* ≤ 0.001, \*\*\*\**p* ≤ 0.0001. Control mRNAs: non-slip (ns), in-frame control (ifc) and wild-type (wt) are coloured in teal. Stem 1 mutants are coloured in dark blue. Stem 2 mutants are coloured in dark purple. C) Diagram of dual luciferase assay used to measure frameshifting efficiency in HEK293T and BHK-21 cells. Cells were co-transfected with two plasmids to transiently express a dual luciferase reporter cassette containing sequences of interest, along with either wild-type or inactive 2A_R85A/R87A_ protein. Elements are colour-coded as in A). D, E) Calculated −1 frameshift efficiencies from luminometry measurements in D) BHK-21 cells 36 hpt and E) HEK293T cells 24 hpt. Data are represented as the mean ± SD error bars, normalised to the in-frame control (ifc) (*n* = 4 biological repeats for BHK-21 and *n* = 3 biological repeats for HEK293T). ns, not significant, \**p* ≤ 0.05, \*\**p* ≤ 0.01, \*\*\**p* ≤ 0.001, \*\*\*\**p* ≤ 0.0001.

Next, we assessed the effects of S1 and S2 mutations on frameshifting in HEK293T and BHK-21 cells. We co-transfected cells with a dual-luciferase reporter plasmid harbouring the TMEV frameshift site, and an expression plasmid for either wild-type 2A or inactive 2A_R85A/R87A_ (**Figure 5C**). In these experiments, firefly luciferase (F-Luc) is produced by a −1 frameshift, and frameshifting efficiency is calculated relative to an in-frame control (ifc) where R-luc and F-Luc are in the same reading frame (**STAR Methods**). Notably, frameshifting efficiency on wild-type RNA was lower than *in vitro* (23.6±4.7% in BHK-21, **Figure 5D** and 17.8±0.9% in HEK293T, **Figure 5E**), possibly due to our inability to control cytoplasmic 2A concentration. Nevertheless, assays with both cell lines yielded similar results (**Figure 5D, 5E**). All S1 mutations were strongly inhibitory to −1 PRF (<1.6%) and the ΔU19 mutation had no residual activity in cells, in contrast to *in vitro* experiments. S2 mutants also displayed similar trends: the stem-swap or shortening by −1 bp were mildly inhibitory to −1 PRF, whereas lengthening by +1 bp or shortening by −2 bp were strongly inhibitory. Overall, our translation assays *in vitro* and in cells support the critical importance of 2A-dependent S1 formation as the primary determinant of ribosomal frameshifting.

### The first base pair of stem 1 exhibits intrinsic plasticity

In many PRF stimulatory pseudoknots, S1 stability is essential for frameshifting — especially of the first base pairs to be encountered by the ribosome^61–63^. Our observation that only the second and third base pairs of S1 are stabilised by the arginine loop (**Figure 2**) was therefore unexpected. We wondered whether single-nucleotide mutations in S1 would prevent rearrangement into the pseudoknot, and designed a series of smFRET reporters to evaluate the relative importance of base pairing at all three positions (**Figure 6A**). Instead of mutating the guanines that interact with R85 and R87, we targeted the cytosines on the opposite strand (C38G, C37G and C36G), with the triple mutant (CCC→GGG) as a control for completely preventing pseudoknot formation. All individual S1 mutations abolished 2A binding by EMSA and MST (**Figure 6B, 6C**) and none could adopt the characteristic mid-FRET state (E = 0.63, **Figure 3E**) corresponding to pseudoknot formation (**Figure 6D**), emphasising the importance of base pairing at all three positions. However, addition of 2A to the C38G mutant produced a minor population of molecules with a pseudoknot-like conformation (E_1_ = 0.76). This was also present, to a lesser extent, in the C36G dataset (E_1_ = 0.78). These could not be detected by ensemble methods, consistent with them being unstable, short-lived intermediates. Nevertheless, this suggests some intrinsic plasticity at the first and last position of S1.

**Figure 6.**
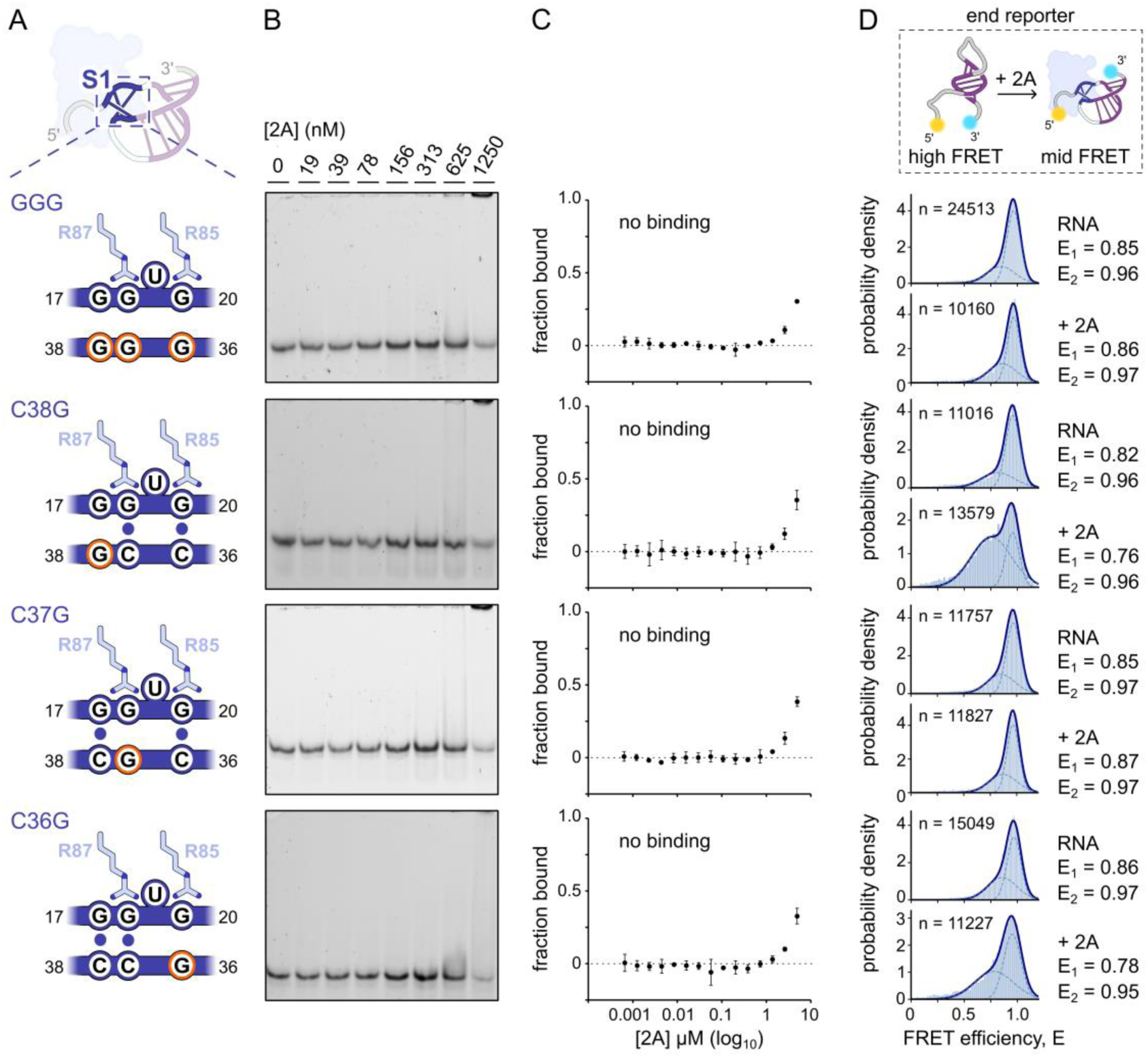
All three base pairs of stem 1 are critical for pseudoknot formation. A) Diagram showing the location of mutated nucleotides in stem 1 (S1) that disrupt base pair formation but not arginine-guanine interactions. B) EMSA analysis of the effects of disrupting individual S1 base pairs on 2A binding. C) MST binding curves and approximate K_D_ values using the same S1 mutant RNAs, assuming 1:1 binding stoichiometry. Experiments were indepedently performed twice, with error bars indicating ± SD. D) Corrected FRET efficiency (E) histograms of S1 mutant end reporter RNAs, combined from three biological repeats. Showing data for (*upper*) RNA in isolation, (lower) RNA in the presence of 0.5 µM wild-type 2A. Corrected average FRET efficiencies (E) and number of molecules (n) are reported in each histogram. Gaussian fits are shown as dashed curves for two Gaussians and a solid blue line for a single Gaussian. R^2^ = 0.99 for all Gaussian fits.

### The 2A-pseudoknot interaction is likely universal in cardioviruses

The S1-forming GGxG and CCC motifs are highly conserved in cardiovirus genomes^49^. Furthermore, although 2A protein sequences are divergent within the genus, a pair of arginines (equivalent to TMEV R85 and R87) are conserved across *Cardiovirus theileri*, *Cardiovirus rueckerti, Cardiovirus saffoldi*, *Cardiovirus dhusarah* and *Cardiovirus rudhira*^52^. To investigate this further, we turned our attention to EMCV (*Cardiovirus rueckerti*), for which the structure of 2A protein is known^53^ but the minimal RNA stimulatory element is 10 nt larger than that of TMEV (**Figure S6A**). Structural alignment of the EMCV 2A *apo* structure to the 2A-pseudoknot complex reported here showed that the RNA binding surface is likely compatible with S1 formation (**Figure S6B**). EMCV 2A arginine loop residues R97 and R95 are juxtaposed with the second and third base pairs of S1 respectively, and the required flexibility for the “G-clamp” is also present: a 5.9 Å movement at glycine 33 (equivalent to TMEV 2A glycine 27) is observed between different chains in the asymmetric unit related by non-crystallographic symmetry.

To test whether the EMCV stimulatory element forms an equivalent pseudoknot upon 2A binding, we designed a doubly-labelled Cy5/Cy3 ‘end reporter’ RNA (**Figure S6C**) and conducted smFRET measurements in the presence and absence of EMCV 2A protein (**Figure S6D**). We observe a major high FRET state in the absence of 2A (E_2_ = 0.98±0.01) which decreases (E = 0.60±0.02) in the presence of 2A. To test whether this change is due to pseudoknot formation, we conducted experiments with a C47G mutant RNA (equivalent to TMEV C36G) to disrupt S1. This rendered the EMCV RNA unresponsive to 2A protein (**Figure S6E**).

By manually enforcing S1 base pairs, we used SimRNA^64^ to predict the structure of the EMCV pseudoknot (**Figure S6F**). As expected, the model has a side-by-side arrangement of S1 (3 bp) and S2 (7 bp), and a flipped-out A19 base in S1. The additional 10 nucleotides in EMCV correspond to an extended L2 sequence, predicted to form a tetraloop-capped 4 bp stem that stacks against S2. The absence of L1 in EMCV, but the introduction of an L3 (U48) alters the angle between S1 and S2, making the helical axes almost parallel. However, manual docking of this predicted RNA onto the *apo* structure of EMCV 2A illustrates that the principles of RNA recognition are identical (**Figure S6G**). Overall, the pseudoknot architecture and its binary switching mechanism are likely conserved across all cardioviruses.

## Discussion

### An unusual RNA pseudoknot architecture

Here we describe a previously unknown class of RNA pseudoknot, the formation of which is strictly conditional upon specific interactions with 2A protein. The inability of AlphaFold3 to predict this complex underscores an urgent need for more experimental structures of diverse RNAs and RNA-protein complexes, to better inform deep learning-based approaches^42,65^.

In the protein-bound state, the TMEV stimulatory element topologically resembles an H-type pseudoknot, with a 3 bp S1 connected to a 7 bp S2. However, our structure highlights major differences to structurally characterised PRF pseudoknots. Many of these possess two coaxially stacked stems (S1 and S2), exemplified by beet western yellow virus (BWYV; *Polerovirus BWYV*)^26,66^, pea enation mosaic virus (PEMV; *Enamovirus PEMV*)^67^, potato leaf roll virus (PLRV; *Polerovirus PLRV*)^28^, sugarcane yellow leaf virus (ScYLV; *Polerovirus SCYLV*)^31^ and SARS-CoV-2 (*Betacoronavirus pandemicum*)^34,35^. In these structures, folding is nucleated by a stable 5′ S1 of at least 5 bp, reinforced by tertiary interactions with nucleotides in loops (reviewed in^58^). In the TMEV RNA, the absence of L3 makes coaxial stacking of S1/S2 impossible: instead they exist in a side-by-side arrangement, with a ∼55° rotation between helical axes, and L2 adenine nucleotides contribute to tertiary interactions with S1.

Why does this pseudoknot depend on 2A protein? In RNA-only experiments, the molecule exists exclusively as a stem-loop (S2), and S1 does not form, even though intuitively the pseudoknot (10 bp, 27 Watson-Crick hydrogen bonds) appears to be more stable than the stem-loop (7 bp, 18 Watson-Crick hydrogen bonds). Our structure suggests that the answer to this question is related to the sub-optimal geometry of S1. When RNA pseudoknots were first described, three base pairs were proposed as a theoretical minimum for S1 stability^68^, and our discovery of a flipped-out base within the 3 bp S1 was unexpected. The energetic penalty associated with displacement and solvation of U19 likely explains the dependency of S1 formation on favourable arginine-guanine interactions. In this way, 2A can be thought of as a catalyst or chaperone, reducing the energy barrier for the stem-loop to pseudoknot transition. Once formed, the pseudoknot is further stabilised by interactions with 2A (8 protein-RNA hydrogen bonds, 634 Å^2^ interface) consistent with the contribution of ΔH (−10.7±0.13 kcal/mol) to the overall ΔG of 2A binding (−9.8 kcal/mol)^52^. This explains the ability of the complex to function as a blockade during ribosome elongation.

### High-efficiency frameshifting in cardioviruses

Why is the TMEV frameshift efficiency so high? Numerous biochemical and virological studies have identified the stability of S1 — in particular, the first base pairs of S1 — as a major determinant of frameshifting^61–63^. A striking feature of the TMEV 2A-RNA complex is that the arginine loop exclusively stabilises the second and third positions of S1, not the first. Furthermore, our smFRET data suggests that transient 2A-RNA interactions may still occur in the absence of the first base pair. We hypothesise that as the ribosome approaches S1, reversible breaking and reforming of the first base pair may occur before 2A dissociates, thereby creating tension on the mRNA that favours a −1 frameshift. This could contribute to the unusually high PRF efficiency (∼85%) in cardioviruses that cannot be explained by thermodynamic equilibrium models^22,23^. Additional evidence for mRNA tension in this system comes from the observation that reducing the spacer length by one nucleotide biases the ribosome towards −2 frameshifting^51^, also observed at the HIV-1 *gag/pol* site^19^.

Our structure also resolves a mystery regarding the unusually long spacer between the PRF slippery sequence and downstream stimulatory element. This was initially reported to be 14 nt, based on bioinformatic identification of S2 and experiments demonstrating its importance for PRF^48,50^. However, our discovery of S1 in the 2A-RNA complex reduces the spacer to 9 nt between the end of the GGUUUUU slippery sequence the start of the GGxG motif comprising the 5′ strand of S1, comparable to other PRF signals^2^. Interestingly, the first contact between 2A and the PRF stimulatory RNA is two nucleotides earlier than S1, with A15 stacking against W89. Although this is not required for 2A binding (**Figure S5**) it may nevertheless contribute to slowing down elongation as the ribosome approaches the first base pair of S1^14^.

### A protein-dependent riboswitch?

In this system, the RNA behaves as a binary switch, converting from stem-loop to pseudoknot in the presence of 2A. In contrast, the 2A protein undergoes minimal conformational change during complex formation (C_α_ backbone RMSD of 0.39 Å). We therefore propose this element represents a protein-dependent riboswitch. Generally, riboswitches modulate gene expression through interconversion between structurally distinct conformations in response to a ligand (reviewed in^58,69,70^). To date, riboswitches have been described for a wide variety of ligands, including nucleotide precursors (pre-queuosine_1_, preQ_1_^71^), metabolites (coenzyme B12^72^, S-adenosylmethionine, SAM^73,74^), amino acids (lysine^75^, glycine^76^, glutamine^77^) and even tRNAs (T-box^78^). In our case, the 2A protein itself acts as the primary ligand. Although this is unusual, ribosomal proteins have been shown to modulate the stability of adenine-sensing riboswitches^79^. Another conceptual difference here is that the TMEV PRF RNA exerts its regulatory effect at the level of translation elongation, rather than translation initiation or transcription termination.

A feature essential to riboswitches is the stabilisation of the ligand-sensing ‘aptamer’ domain, typically S2, by the cognate ligand. This is common in bacterial riboswitches that regulate translation initiation, as the Shine Dalgarno sequences are occluded within the stabilised S2. In our case the ‘aptamer’ would be the GGxG motif comprising S1, and the ligand would be the arginine loop of 2A. The selectivity for arginine is demonstrated by the complete inability of the 2A_R85A/R87A_ protein to induce switching (**Figure 3**) and no evidence for spontaneous sampling of the pseudoknot conformation in the absence of protein. This contrasts with the preQ_1_ riboswitch, which transiently samples an intrinsically unstable pseudoknot-like fold in the absence of preQ_1_ ligands^80^.

In conclusion, this work defines the molecular basis for the highest efficiency viral frameshifting event in nature. Reframing the idea of a PRF stimulatory element as a riboswitch has vast implications. We speculate that other canonical PRF elements — especially those reported to exhibit multiple conformations — might also be regulated by cellular proteins, viral proteins or metabolites. Indeed, ribosome profiling of cells infected with mouse hepatitis virus (MHV; *Betacoronavirus muris*^81^), SARS-CoV-2^82,83^ and PRRSV^84^ has revealed variations in frameshifting over time at PRF signals previously thought to function at a fixed efficiency. Understanding this hidden layer of regulation will be essential to realise the potential of artificial intelligence in designing ligands to manipulate these elements^85,86^.

### Limitations of the Study

While the SAXS and FRET data support the presence of a single RNA conformation in the absence of 2A, the possibility of transient pseudoknot sampling under different conditions (ionic strength, temperature, molecular crowding, or in the presence of the ribosome) cannot be excluded. Differences in gaussian peak widths observed in our smFRET measurements could reflect either a broader ensemble of RNA conformations, enhanced flexibility in the 5′ or 3′ ends and/or increased dynamic interconversion between states occurring on timescales faster than the measurement resolution. Our current data do not allow us to unambiguously distinguish between static and dynamic contributions.

## Resource Avalibility

### LEAD CONTACT

- Requests for further information and resources should be directed to and will be fulfilled by the lead contact, Chris H. Hill.

### MATERIALS AVAILBILITY

- Plasmid maps generated in this study have been deposited to Mendeley Data **10.17632/78kj389b6v.1**.

### DATA AND CODE AVAILBILITY

- Crystallographic data and refined atomic models for the 2A-RNA complex have been deposited to the Worldwide Protein Data Bank (wwPDB) under accession code 9RVP. Small-angle X-ray scattering data has been deposited to the Small Angle Scattering Biological Data Bank (SASBDB) under accession codes SASDWQ5 (2A protein only), SASDWR5 (RNA only) and SASDWS5 (2A-RNA complex). Original gel images, smFRET raw hdf5 data files and Jupyter Notebooks used for smFRET data analysis and have been deposited at Mendeley Data **10.17632/78kj389b6v.1**. These data are publicly available as of the date of publication.
- This paper does not report original code.
- Any additional information required to reanalyze the data reported in this paper is available from the lead contact upon request.

## Acknowledgments

We acknowledge the YSBL X-ray crystallography facility supported by BBSRC (BB/T017805/1), Diamond Light Source for access to beamline I04 under proposal mx-32736, and EMBL for access to the bioSAXS beamline P12 under proposal SAXS-1125. We thank Johan Turkenburg and Sam Hart for technical assistance at the YSBL X-ray crystallography facility. Remote synchrotron access was supported in part by the EU FP7 infrastructure grant BIOSTRUCT-X (Contract No. 283570). We thank the Molecular Interactions team (Andrew Leech, Katy Cornish and Alex Payne-Dwyer) and the Protein Production team (Jared Cartwright and Matt Tyreman) for technical support within the York Bioscience Technology Facility. We also thank Ben Butt for helpful discussions. J.K.B. is supported by a Medical Research Council DiMeN PhD studentship (MR/W006944/1). C.M.J. acknowledges the European Union’s Horizon 2020 research and innovation programme “iNEXT Discovery” grant agreement no. 871037. M.A.S.A was supported by BBSRC grant to TDC (BB/T008032/). T.D.C. is supported by BBSRC (BB/W00061X/1). M.C.L. is supported by BBSRC (BB/W000555/1) and EPSRC (EP/Y000501/1). S.D.Q. is supported by Alzheimer’s Research UK (RF2019A-001) and EPSRC (EP/X525856/1, EP/V034030/1). S.P.G. was supported by an EPSRC studentship (EP/T518025/1). H.C.Y.K. was funded by an Exciting Instruments (Sheffield, UK) studentship administered by Generation Research (University of York, UK). S.C.G. was funded by a Sir Henry Dale fellowship (098406/Z/12/B) from the Wellcome Trust and the Royal Society. We acknowledge BBSRC funding to I.B. and C.H.H. (BB/V000306/1). T.C.P and C.H.H. are supported by a Sir Henry Dale Fellowship (221818/Z/20/Z) from the Wellcome Trust and the Royal Society (to C.H.H). C.H.H. is supported by the Lister Institute of Preventive Medicine. For the purpose of open access, the authors have applied a CC BY public copyright licence to any Author Accepted Manuscript version arising from this submission.

## Author Contributions

Conceptualization, J.K.B., C.H.H. Data Curation, J.K.B., C.M.J., C.H.H. Formal Analysis, J.K.B., C.M.J., T.C.P., C.H.H. Funding Acquisition, S.C.G, C.H.H. Investigation, J.K.B., C.M.J., T.C.P., H.C.Y.K., S.P.G., I.B., C.H.H. Methodology, J.K.B., C.M.J., M.A.S.A., Project Administration, S.D.Q., C.H.H. Resources, J.K.B., C.M.J., I.B., C.H.H. Software, M.A.S.A., T.D.C., Supervision, M.A.S.A., J.A.L.H., M.C.L., S.D.Q., C.H.H. Validation, J.K.B., C.M.J., C.H.H. Visualization, J.K.B., C.H.H. Writing – original draft, J.K.B., C.H.H., Writing – review & editing, J.K.B., T.D.C., S.C.G., I.B, M.C.L., S.D.Q., C.H.H.

## Declaration of Intrests

T.D.C. and M.A.S.A. are employees and shareholders in Exciting Instruments Ltd.

## STAR Methods

### KEY RESOURCES TABLE

| REAGENT or RESOURCE | SOURCE | IDENTIFIER |
| --- | --- | --- |
| Bacterial and virus strains |  |  |
| *Escherichia coli* Stellar cells | Takara | Cat# 636763 |
| *Escherichia coli* BL21(DE3) pLysS cells | Sigma | Cat# 69451-3 |
| Chemicals, peptides, and recombinant proteins |  |  |
| 2 × TY broth capsules | Melford | Cat# X15640 |
| DMEM | Gibco | Cat# 41966-029 |
| Fetal Bovine Serum | Gibco | Cat# A5256701 |
| Penicillin-Streptomycin (5,000 U/mL) | Gibco | Cat# 15070063 |
| L-Glutamine (200 mM) | Gibco | Cat# 25030149 |
| Isopropyl-β-D-thio-galactopyranoside (IPTG) | Melford | Cat# I56000 |
| Trizma® base | Sigma | Cat# T1503 |
| Sodium chloride | Fisher Chemical | Cat# S/3161/65 |
| Imidazole | Sigma | Cat# 56750 |
| DNase I | Sigma | Cat# D4527 |
| DTT | Melford | Cat# D11000 |
| Potassium chloride | Sigma | Cat# P9541 |
| Glycerol | Honeywell | Cat# 49770-2.5L |
| HEPES | Fluorochem | Cat# F047861 |
| Ammonium acetate | Sigma | Cat# A1542 |
| Bis-Tris | Sigma | Cat# B9754 |
| Poly(ethylene glycol) 3350 | Sigma | Cat# 88276 |
| Magnesium acetate tetrahydrate | Fluorochem | Cat# F449646 |
| Bromophenol Blue sodium salt | Sigma | Cat# B5525 |
| ProtoGel (30%) | National Diagnostics | Cat# EC-890 |
| AccuGel 19:1 (40%) | National Diagnostics | Cat# EC-850 |
| Urea, UltraPure | Melford | Cat# U20200 |
| TEMED | Melford | Cat# T18000 |
| Ammonium persulfate (APS) | Melford | Cat# A20500 |
| Bovine liver tRNA | Sigma | Cat# R4752 |
| UltraPure BSA (50 mg/mL) | Invitrogen | Cat# 10743447 |
| Magnesium chloride, hexahydrate | Millipore | Cat# 442611-M |
| AzDye647-maleimide | Vector Labs | Cat# FP-1122-1 |
| Cy5-maleimide | GE Healthcare | Cat# PA25031 |
| NdeI | New England Biolabs | Cat# R0111S |
| dNTP bundle | Jena Bioscience | Cat# NU-1005S |
| RiboLock RNase inhibitor | Thermo Fisher | Cat# EO0381 |
| L-[S35] Methionine 10 mCi/ml. Translation grade. | Hartmann Analytic | Cat# SRM-01 |
| TransIT®-LT1 transfection reagent | Mirus Bio | Cat# MIR 2304 |
| PEI transfection reagent | MedChemExpress | Cat# HY-K2014 |
| Passive Lysis 5X buffer | Promega | Cat# E1941 |
| Recombinant protein: wild type TMEV 2A | This paper, based on Hill *et al.*<sup>52</sup> | N/A |
| Recombinant protein: TMEV inactive 2A<sub>R85A/R87A</sub> | This paper, based on Hill *et al.*<sup>52</sup> | N/A |
| Recombinant protein: TMEV 2A<sub>R7A</sub> | This paper | N/A |
| Recombinant protein: TMEV 2A<sub>K24A</sub> | This paper | N/A |
| Recombinant protein: TMEV 2A<sub>H26A</sub> | This paper | N/A |
| Recombinant protein: TMEV 2A<sub>W89A</sub> | This paper | N/A |
| Recombinant protein: EMCV wild type 2A | This paper, based on the work of Hill *et al.*<sup>53</sup> | N/A |
| Critical commercial assays |  |  |
| 5' EndTag DNA/RNA Labeling Kit | Vector Labs | Cat# MB-9001 |
| Flexi® Rabbit Reticulocyte Lysate System | Promega | Cat# L4540 |
| Dual-Glo® Luciferase Assay System | Promega | Cat# E2920 |
| Dual-Luciferase® Reporter Assay System | Promega | Cat# E1910 |
| Deposited data |  |  |
| Raw data files | This paper | Mendeley data: 10.17632/78kj389b6v.1 |
| Crystal structure of the 2A-RNA pseudoknot complex | This paper | PDB: 9RVP |
| Small angle scattering data: 2A protein | This paper | SASBDB: SASDWQ5 |
| Small angle scattering data: RNA | This paper | SASBDB: SASDWR5 |
| Small angle scattering data: 2A-RNA complex | This paper | SASBDB: SASDWS5 |
| Crystal structure of TMEV 2A protein | Hill *et al.*<sup>52</sup> | PDB: 7NBV |
| Crystal structure of EMCV 2A protein | Hill *et al.*<sup>53</sup> | PDB: 7BNY |
| Experimental models: Cell lines |  |  |
| BHK-21 | ATCC | CCL-10 |
| HEK293T | ATCC | CRL-3216 |
| Oligonucleotides |  |  |
| See Table S1 for DNA oligonucleotides | This paper | N/A |
| See Table S2 for RNA oligonucleotides | This paper | N/A |
| Recombinant DNA |  |  |
| pGEX6P2 vector containing wild type TMEV 2A (used for subcloning) | Napthine *et al.*<sup>51</sup> | CH plasmid #73 |
| pOPTnH vector containing inactive TMEV 2A<sub>R85A/R87A</sub> (used for subcloning) | Hill *et al.*<sup>52</sup> | CH plasmid #33 |
| pOPTHMBP TMEV wt 2A (used to make protein) | This paper | CH plasmid #219 |
| pOPTHMBP TMEV 2A<sub>R85A/R87A</sub> (used to make protein) | This paper | CH plasmid #423 |
| pOPTHMBP TMEV 2A<sub>R7A</sub> (used to make protein) | This paper | CH plasmid #323 |
| pOPTHMBP TMEV 2A<sub>K24A</sub> (used to make protein) | This paper | CH plasmid #324 |
| pOPTHMBP TMEV 2A<sub>H26A</sub> (used to make protein) | This paper | CH plasmid #325 |
| pOPTHMBP TMEV 2A<sub>W89A</sub> (used to make protein) | This paper | CH plasmid #326 |
| pOPTnH EMCV wt 2A (used to make protein) | Hill *et al.*<sup>53</sup> | CH plasmid #31 |
| pUCGW_TMEV-ns-SUMO (non-slip control for RRL *in vitro* assays) | This paper | CH plasmid #433 |
| pUCGW_TMEV-ifc_SUMO (in-frame control for RRL *in vitro* assays) | This paper | CH plasmid #434 |
| pUCGW_TMEV-wt_SUMO (wild type sequence for RRL *in vitro* assays) | This paper | CH plasmid #432 |
| pUCGW_TMEV-S1-swap_SUMO (S1 swap test construct for RRL *in vitro* assays) | This paper | CH plasmid #428 |
| pUCGW_TMEV-S1-AUstem_SUMO (S1 AU stem test construct for RRL *in vitro* assays) | This paper | CH plasmid #429 |
| pUCGW_TMEV-S1-UAstem_SUMO (S1 UA stem test construct for RRL *in vitro* assays) | This paper | CH plasmid #430 |
| pUCGW_TMEV-S1-deltaU19_SUMO (S1 ΔU19 test construct for RRL *in vitro* assays) | This paper | CH plasmid #431 |
| pUCGW_TMEV-S2-plus1bp_SUMO<br>(S2 +1 bp test construct for RRL *in vitro* assays) | This paper | CH plasmid #424 |
| pUCGW_TMEV-S2-minus1bp_SUMO<br>(S2 -1 bp test construct for RRL *in vitro* assays) | This paper | CH plasmid #425 |
| pUCGW_TMEV-S2-minus2bp_SUMO<br>(S2 -2 bp test construct for RRL *in vitro* assays) | This paper | CH plasmid #426 |
| pUCGW_TMEV-S2-swap_SUMO<br>(S2 swap test construct for RRL *in vitro* assays) | This paper | CH plasmid #427 |
| pSGDlucV3.0 | Gifted from Betty Chung<sup>121</sup> | Loughran et al.<sup>120</sup><br>addgene #119760 |
| pSGDluc_TMEV-wt<br>(wild type sequence for *vivo* dual luciferase assays) | This paper | CH plasmid #396 |
| pSGDluc_TMEV-ifc<br>(in-frame control for *in vivo* dual luciferase assays) | This paper | CH plasmid #397 |
| pSGDluc_TMEV-S2-plus1bp<br>(S2 + 1 bp for *in vivo* dual luciferase assays) | This paper | CH plasmid #398 |
| pSGDluc_TMEV-S2-minus1bp<br>(S2 - 1 bp for *in vivo* dual luciferase assays) | This paper | CH plasmid #399 |
| pSGDluc_TMEV-S2-minus2bp<br>(S2 - 2 bp for *in vivo* dual luciferase assays) | This paper | CH plasmid #400 |
| pSGDluc_TMEV-S2-swap<br>(S2 swap for *in vivo* dual luciferase assays) | This paper | CH plasmid #401 |
| pSGDluc_TMEV-S1-swap<br>(S1 swap for *in vivo* dual luciferase assays) | This paper | CH plasmid #402 |
| pSGDluc_TMEV-S1-AUstem<br>(S1 AU stem for *in vivo* dual luciferase assays) | This paper | CH plasmid #403 |
| pSGDluc_TMEV-S1-UAstem<br>(S1 UA stem for *in vivo* dual luciferase assays) | This paper | CH plasmid #404 |
| pSGDluc_TMEV-S1-deltaU19<br>(S1 ΔU19 for *in vivo* dual luciferase assays) | This paper | CH plasmid #405 |
| pCEP4 mammalian expression vector | Invitrogen | Cat# V04450 |
| pCEP4-TMEV-2A-wt<br>(for transient expression of wild type 2A *in vivo* dual luciferase assays) | This paper | CH plasmid #406 |
| pCEP4-TMEV-2A<sub>R85A/R87A</sub><br>(for transient expression of inactive 2A<sub>R85A/R87A</sub> *in vivo* dual luciferase assays) | This paper | CH plasmid #407 |
| Software and algorithms |  |  |
| XIA2 automated pipeline | Winter<sup>89</sup> | https://xia2.github.io/index.html |
| XDS | Kabsch<sup>90</sup> | https://xds.mr.mpg.de/ |
| AIMLESS | Evans and Murshudov<sup>91</sup> | https://www.ccp4.ac.uk/ |
| PHASER | McCoy *et al.*<sup>93</sup> | https://www.ccp4.ac.uk/ |
| AlphaFold3 | Abramson *et al.*<sup>57</sup> | https://alphafoldserver.com |
| COOT | Emsley *et al.*<sup>94</sup> | https://www2.mrc-lmb.cam.ac.uk/personal/pemsley/coot/ |
| phenix.refine (v1.20.1) | Liebschner *et al.*<sup>95</sup> | https://www.phenix-online.org/ |
| MolProbity | Chen *et al.*<sup>96</sup> | https://www.phenix-online.org/ |
| PyMol | Schrödinger, LLC<sup>97</sup> | https://www.pymol.org/ |
| ChimeraX | Meng *et al.*<sup>98</sup> | https://www.cgl.ucsf.edu/chimerax/ |
| 3DNA webserver | Li *et al.*<sup>99</sup> | https://web.x3dna-dssr.org/ |
| ATSAS software suite (v3.2.1) | Manalastas-Cantos *et al.*<sup>103</sup> | https://www.embl-hamburg.de/biosaxs/software.html |
| CHROMIXS | Panjikovich *et al.*<sup>101</sup> | https://www.embl-hamburg.de/biosaxs/software.html |
| SASFLOW pipeline | Franke *et al.*<sup>102</sup> | https://www.embl-hamburg.de/biosaxs/software.html |
| Correlation Map (CorMap) | Franke *et al.*<sup>103</sup> | https://www.embl-hamburg.de/biosaxs/software.html |
| Primus | Manalastas-Cantos *et al.*<sup>104</sup> | https://www.embl-hamburg.de/biosaxs/software.html |
| *datvc* | Rambo and Tainer<sup>106</sup> | https://www.embl-hamburg.de/biosaxs/software.html |
| *datmow* module | Fischer *et al.*<sup>107</sup> | https://www.embl-hamburg.de/biosaxs/software.html |
| GNOM (v4.6) | Svergun<sup>109</sup> | https://www.embl-hamburg.de/biosaxs/software.html |
| MONSA (v1.44) | Svergun<sup>110</sup> | https://www.embl-hamburg.de/biosaxs/software.html |
| MULCh | Whitten *et al.*<sup>111</sup> | https://smb-research.smb.usyd.edu.au/NCVWeb/ |
| DAMMIN (v5.3) | Svergun<sup>110</sup> | https://www.embl-hamburg.de/biosaxs/software.html |
| SREFLEX | Panjikovich & Svergun<sup>113</sup> | https://www.embl-hamburg.de/biosaxs/software.html |
| CORAL (v1.1) | Petoukhov *et al.*<sup>105</sup> | https://www.embl-hamburg.de/biosaxs/software.html |
| CRY SOL (v2.8.3) | Svergun *et al.*<sup>114</sup> | https://www.embl-hamburg.de/biosaxs/software.html |
| Anaconda Navigator | Anaconda | https://anaconda.org/ |
| Jupyter Notebooks (v6.5.4) | Jupyter | https://jupyter.org/ |
| FRET Bursts (v0.8.3) | Ingargiola *et al.*<sup>116</sup> | https://fretbursts.readthedocs.io/en/latest/# |
| FRET-restrained positioning and screening (FPS) tool (v1.1) | Kalinin *et al.*<sup>119</sup> | https://www.mpc.hhu.de/software/fps |
| MO.Affinity Analysis software | NanoTemper Technologies | Cat# MO-S001A |
| ImageStudio 6.1 | LICORbio | https://www.licorbio.com/image-studio |
| Spyder (v5.4.3) | Spyder IDE | https://docs.spyder-ide.org/current/index.html |
| SciPy (v1.10.1) | SciPy | https://docs.scipy.org/doc/scipy-1.10.1/index.html |
| InkScape (v1.4.2) | Inkscape | https://inkscape.org/ |

| Other |  |  |
| --- | --- | --- |
| Ni Sepharose™ 6 Fast Flow histidine-tagged protein purification resin | Cytiva | Cat# 17531802 |
| HiTrap™ Heparin HP affinity columns | Cytiva | Cat# 17040703 |
| HiLoad™ 16/600 Superdex 75 pg | Cytiva | Cat# 28989333 |
| SnakeSkin™ dialysis tubing 10K MWCO | Thermo Fisher | Cat# 88243 |
| Amicon® Ultra Centrifugal Filter, 10 kDa MWCO | Merck | Cat# UFC901024 |
| 4–20% Mini-PROTEAN® TGX™ Precast Protein Gels | Bio Rad | Cat# 4561093 |
| Protein LoBind® Tubes | Eppendorf | Cat# 0030108116 |
| DNA LoBind® Tubes | Eppendorf | Cat# 0030108051 |
| Monolith Premium Capillaries | NanoTemper | Cat# MO-K025 |
| Cover slips, round #1, 30 mm | VWR | Cat# 631-0174 |
| RNA Clean & Concentrator-5 | Zymo Research | Cat# R1015 |

## EXPERIMENTAL MODEL AND STUDY PARTICIPANT DETAILS

### Bacterial culture

All gene cloning, manipulation and plasmid propagation steps were performed in *Escherichia coli* Stellar cells (Takara) cultured in LB media with appropriate selection antibiotics.

Recombinant proteins: wild-type 2A, inactive 2A_R85A/R87A_, 2A_R7A_, 2A_K24A_, 2A_H26A_ and 2A_W89A_ were expressed in *E.coli* BL21(DE3) pLysS cells grown in 2x TY media supplemented with appropriate antibiotics at 37°C, 200 rpm, until an optical density at 600 nm (OD_600_) of 0.5 - 0.7 was reached. Expression was induced with 1.0 mM isopropyl β-d-1-thiogalactopyranoside (IPTG) and continued overnight (200 rpm, 21°C, 16 h). Bacteria were harvested (4000 × g, 4°C, 20 min), washed in cold 20 mM Tris-HCl pH 8.0 and stored at −20°C.

### Cell lines

HEK293T cells (ATCC) were maintained in Dulbecco’s modified Eagle’s medium (DMEM) supplemented with 10% fetal bovine serum (FBS) at 37°C and 5% CO_2_. BHK-21 cells (ATCC) were maintained in DMEM supplemented with 10% fetal bovine serum (FBS), 100 U/mL penicillin, 100 µg/mL streptomycin and 2 mM L-glutamine at 37°C and 5% CO_2_.

## METHOD DETAILS

### Cloning

TMEV 2A cDNA was amplified by PCR from plasmid pOPT3G-based constructs (previously described in^52^) and cloned into pOPTHMBP to introduce a TEV protease-cleavable N-terminal His_6_ MBP tag. Mutant TMEV 2A proteins were prepared by mutagenesis PCR using the primers detailed in **Table S1**.

For *in vitro* translation assays, we designed reporter constructs with an EMCV internal ribosome entry site (IRES) to drive translation of a polypeptide including a 3x Myc tag, followed by nucleotides 4137-4307 of the TMEV genome (GenBank: NC_001366.1, corresponding to 35 codons upstream of the frameshift site, the slippery sequence and the stimulatory element), followed by the coding sequence for SUMO protein in the −1 frame. Constructs were designed such that if elongation continues in the 0 frame the product is ∼19 kDa, whereas shifting into the −1 frame enables translation to the end of SUMO, yielding a ∼28 kDa product. To enable this, minor modifications were made to the wild-type TMEV sequence, as follows. The StopGo signal immediately before the frameshift site was inactivated by mutation of the NPGP motif to NPLV (insertion of CT between nucleotides 4229 and 4230, and deletion of C4234 and T4235). A −1 frame stop codon (UAA) within the stimulatory element was removed by introducing the T4274A mutation. All further S1 and S2 mutations were made in this context. The non-slippery (ns) control mRNA contains a mutated slippery sequence to prevent shifting into the −1 frame, and the in-frame control (ifc) mRNA introduces another nucleotide after the stimulatory element, to simulate the effect of 100% −1 frameshifting (i.e. by bringing the SUMO coding sequence into the 0 frame). These sequences were synthesised *de novo* and subcloned into the pUC-GW-Amp vector (GENEWIZ, Azenta Life Sciences).

To test frameshifting in cells, DNA sequences (nucleotides 4137-4307 of the TMEV genome, described above) corresponding to the modified wild-type sequence, mutants and controls, were amplified by PCR using primers in **Table S1** and subcloned into the pSGDlucV3.0^120^ dual luciferase reporter plasmid (a gift from Betty Chung^121^) by restriction digest and ligation at the *PspXI*/*BglII* sites, to introduce sequences of interest between an upstream *Renilla* luciferase (R-luc) in the 0 frame, and a downstream Firefly luciferase (F-luc) in the −1 frame. F-Luc is only produced by a −1 frameshift, and the in-frame control (ifc) construct introduces another nucleotide after the stimulatory element, to simulate the effect of 100% −1 frameshifting (i.e. by bringing the F-luc coding sequence into the 0 frame).

Wild-type and inactive 2A_R85A/R87A_ were cloned into the pCEP4 (Invitrogen) mammalian expression vector by restriction digest and ligation at the *XhoI/BamHI* sites. All constructs were verified by sanger sequencing.

### Protein purification

TMEV 2A protein purification was performed as previously described^52^. Cell pellets from 2.0 L of *E.coli* BL21(DE3) pLysS cells were thawed in 300 mL of lysis buffer (50 mM Tris-HCl pH 8.0, 300 mM NaCl, 10 mM imidazole supplemented with 50 µg/mL DNase I and EDTA-free proteases) and lysed using a cell disruptor (Constant Systems, 27 kPSI, 4°C). The insoluble fraction was removed by centrifugation (39,000 × g, 40 min, 4.0°C) and discarded. Supernatant was incubated (1 h, 4°C) with 4.0 mL Ni Sepharose 6 Fast Flow resin (Cytiva) pre-equilibrated in the same buffer. The resin was washed three times with 200 mL buffer (50 mM Tris-HCl pH 8.0, 300 mM NaCl, 10 mM imidazole) in a gravity column. The protein was batch eluted with 20 mL of 50 mM Tris-HCl pH 8.0, 300 mM NaCl, 300 mM imidazole. His_6_-tagged TEV protease (50 µg/mL, prepared in-house) was added to the eluate and was dialysed (3.5K MWCO, 4°C, 16 h) against 2.0 L buffer (20 mM Tris-HCl pH 7.4, 400 mM NaCl, 1.0 mM DTT). Dialysed proteins were subject to heparin-affinity chromatography to remove nucleic acids and the His_6_-MBP solubility tag. Samples were loaded onto a 10 mL HiTrap Heparin column (Cytiva) at 1 mL/min, washed twice with buffer A (20 mM Tris-HCl pH 7.4, 400 mM NaCl, 1.0 mM DTT) and eluted with a 0% → 50% gradient of buffer B (20 mM Tris-HCl pH 7.4, 1.0 M NaCl, 1.0 mM DTT) over 8 CV before stepping to 100% buffer B over 10 CV. Fractions containing the 2A protein were pooled and concentrated in an Amicon Ultra centrifugal filter unit (10K MWCO, 4000 × g) prior to size exclusion chromatography (Superdex 75 16/600, 20 mM Tris-HCl pH 7.4, 1.0 M NaCl, 1.0 mM DTT). Purity was assessed by 4-20% gradient SDS-PAGE. Purified protein was either used immediately or was concentrated (∼3.5 mg/mL, 220 µM), snap-frozen in liquid nitrogen and stored at −80°C.

EMCV 2A protein purification was performed as previously described^53^ with minor modifications. Cell pellets from 2.0 L of *E.coli* BL21(DE3) pLysS cells were thawed in 300 mL of lysis buffer (50 mM Tris-HCl pH 8.0, 500 mM NaCl, 30 mM imidazole pH 8.0 supplemented with 50 µg/mL DNase I and EDTA-free proteases) and lysed using a cell disruptor (Constant Systems, 27 kPSI, 4°C). The insoluble fraction was removed by centrifugation (39,000 × g, 40 min, 4.0°C) and discarded. Supernatant was incubated (1 h, 4°C) with 4.0 mL Ni Sepharose 6 Fast Flow resin (Cytiva) pre-equilibrated in the same buffer. The resin was washed five times with 200 mL buffer (50 mM Tris-HCl pH 8.0, 500 mM NaCl, 30 mM imidazole pH 8.0) in a gravity column. The protein was batch eluted with 20 mL of 50 mM Tris-HCl pH 8.0, 500 mM NaCl, 300 mM imidazole and dialysed (3.5K MWCO, 4°C, 16 h) against 2.0 L buffer (50 mM Tris-HCl pH 8.0, 1.0 M NaCl, 1.0 mM DTT). EMCV 2A protein was concentrated in an Amicon Ultra centrifugal filter unit (10K MWCO, 4000 × g) prior to size exclusion chromatography (Superdex 75 16/600, 10 mM Tris-HCl pH 7.4, 1.0 M NaCl, 1.0 mM DTT). Purity was assessed by 4-20% gradient SDS-PAGE. Purified protein was concentrated (∼4 mg/mL, 225 µM), snap-frozen in liquid nitrogen and stored at −80°C.

### 2A-RNA complex formation and crystallisation

RNA oligonucleotides (oligo 1, detailed in **Table S2**) were reconstituted in 10 mM Tris-HCl pH 7.4, 500 mM NaCl to 2.2 mg/mL (200 µM) determined by measuring absorbance at 260 nm using a Nanodrop spectrophotometer (Thermo). 2A-RNA complex was formed by diluting 2A protein to a final concentration of 500 mM NaCl and adding a 1:1 molar ratio of RNA. Complexes were incubated at room temperature for 20 min prior to buffer exchanging into 10 mM HEPES pH 7.4, 100 mM NaCl, 1.0 mM MgCl_2_, 1.0 mM DTT and concentrating using an Amicon Ultra centrifugal filter unit (10K MWCO, 4000 × g, 10°C). Complexes were then subjected to size exclusion chromatography using Superdex 75 16/600 (Cytiva) at 10°C in 10 mM HEPES pH 7.4, 100 mM NaCl, 1.0 mM MgCl_2_, 1.0 mM DTT. Complex formation was assessed by SDS-PAGE and urea-PAGE. Fractions containing the 2A-RNA complex were pooled and concentrated (∼7.5 mg/mL, 693 µM) as assessed by absorbance at 260 nm. Sitting-drop vapor diffusion experiments were set-up in 96-well MRC plates with 50 µL reservoir solution, 150 nL complex and 150 nL crystallisation buffer at 21°C. Diffraction quality crystals grew in 0.2 M ammonium acetate, 0.1 M Bis-Tris pH 5.9, 19% (w/v) PEG 3350. Ten days after plating, crystals were cryo-protected by the addition of 0.2 µL crystallisation buffer supplemented with 20% (v/v) glycerol, prior to harvesting in nylon loops and flash-cooling by plunging into liquid nitrogen.

### X-ray data collection, structure determination, refinement and analysis

Datasets of 3600 images were recorded at Diamond Light Source, beamline I04 (λ = 0.9537 Å) on an Eiger2 XE 16M detector (Dectris), using 100% transmission, an oscillation range of 0.10° and an exposure time of 0.02s per image. Data were processed with XIA2 automated pipeline^89^ using XDS^90^ for indexing and integration and AIMLESS^91^ for scaling and merging. Processing and refinement statistics are detailed in **Table 1**. Resolution cut-off was decided by CC_1/2_ values ≥ 0.5 and an I/σ(I) ≥ 2 in the highest resolution shell^92^. Phasing was done by molecular replacement in PHASER^93^ using two sequential search models. First a partial solution was obtained by placing the coordinates of the *apo* TMEV 2A protein (PDB: <u>7NBV</u>), followed by searching the unit cell for an idealised 7 bp A-form RNA helix (corresponding to S2) generated by AlphaFold3^57^. The model was completed manually by iterative cycles of model building using COOT^94^ and refinement with phenix.refine^95^. MolProbity^96^ was used to assess model geometry through the refinement process. Structural figures (cartoon, stick and surface representations) were rendered in PyMOL (Schrödinger, LLC)^97^ and ChimeraX^98^. Analysis of the RNA pseudoknot geometry was performed using the 3DNA webserver^99^.

### SEC-SAXS

All small angle X-ray scattering (SAXS) experiments were performed coupled to size-exclusion chromatography (SEC-SAXS) at the EMBL P12 beamline at PETRA III (DESY; Hamburg, Germany)^100^. Scattering data were collected on a Pilatus 6M detector at a sample-detector distance of 3 m and at a wavelength of λ = 0.123983 nm (I(s) vs s, where s = 4πsinθ/λ, and 2θ is the scattering angle). For 2A protein-only samples, it was necessary to use high salt to prevent aggregation. 50–100 µL protein at 4.0 mg/ml in 20 mM HEPES pH 7.4, 1.0 M KCl, 2.0% v/v glycerol, 1.0 mM DTT was injected at a 0.35 ml/min flow rate onto a GE Superdex 75 Increase 5/150 column at 20°C. 143 successive 0.25 second frames were collected through two consecutive SEC-SAXS runs, spanning the sample elution peaks of the protein. For RNA-only samples, 50 μl at 7.1 mg/mL in 20 mM HEPES pH 7.4, 400 mM KCl, 2.0% v/v glycerol, 1.0 mM DTT was injected as above; 57 successive 0.25 second frames were collected through the SEC-elution peak. The 2A-RNA complex was prepared at 8.0 mg/mL in an identical buffer and injected as above; data were collected over 41 successive 0.25 second frames. Further details are provided in **Table 2**.

Initial data processing was carried out using CHROMIXS^101^ and azimuthal averaging from 2D to 1D was performed via the SASFLOW pipeline^102^. Buffer frames were evaluated for statistical similarity using all pairwise Correlation Map (CorMap)^103^ p-values, applying a significance threshold (α) of 0.01. Statistically equivalent frames were then averaged to produce the final buffer scattering profile, which was subtracted from the corresponding macromolecular elution peaks. Subtracted data blocks that showed a consistent radius of gyration (*R*_g_) values across the elution profile — assessed via the Guinier approximation — were scaled, compared using CorMap, and averaged to yield the final reduced 1D scattering profiles. All CorMap analyses were conducted using PrimusQT within the ATSAS software suite^104^. Molecular weight estimations were derived using multiple approaches available in ATSAS, including the Porod volume via *datporod*^105^, volume-of-correlation analysis via *datvc*^106^, the *datmow* module^107^ and the Bayesian consensus method^108^. Indirect inverse Fourier transform of the SAXS data was performed using GNOM^109^ to compute the real-space pair-distance distribution function, *p(r)* versus *r*, from which the radius of gyration (*R_g_*) and maximum particle dimension (*D_max_*) were obtained. Further details are provided in **Table 2**.

*Ab initio* dummy atom bead modelling/shape reconstruction was performed using MONSA^110^, that takes into account differences in the average X-ray scattering length density of protein compared to RNA. Briefly, the 2A protein and RNA-only samples were modelled to their individual scattering profiles as single-phase particles, where the average excess X-ray scattering contrast of each phase was set to 2.413 x10^10^ cm^-2^ (2A protein) or 5.580 x10^10^ cm^-2^ (RNA) calculated relative to the average scattering length density of the respective sample buffers using MULCh^111^. Target phase volumes were set to 21.4 nm^3^ (2A protein) and 11.2 nm^3^ (RNA), respectively, based on the amino acid or RNA composition^111^. The 2A-RNA complex was modelled as a two-phase particle against the 2A protein and 2A-RNA complex SAXS datasets in parallel, using the scattering contrasts and target volumes described above for the protein and RNA phases. MONSA was run five times for each particle reconstruction to qualitatively evaluate shape/volume consistency. Additional single-phase dummy atom modelling for the 2A protein or the 2A-RNA complex was performed using DAMMIN^110^ using the output from GNOM as input. Following standard protocols^112^ DAMMIN was run 10 times, the individual models spatially aligned and averaged to develop a consensus shape reconstruction for either sample.

Atomistic models for the RNA were developed using a combination of AlphaFold 3 (AF3)^57^ and normal-mode analysis. The RNA sequence was used as input into AF3 to generate five predicted RNA structures. These individual models underwent subsequent SREFLEX^113^ normal mode analysis, where the RNA structures were divided up into three rigid bodies that underwent repositioning relative to each other minimizing the fit to the RNA SAXS data (nucleotide ranges: 5′-CCACAA-3′; 5′-GGUGC-3′; 5′-GGUGCUAACUAAAUCCCUAGCACCCC-3′). Of the five initial AF3 models it was found that AF3 model 4 (AF3-4) acted as the best template with respect to generating a fit to the SAXS data after SREFLEX analysis. The post SREFLEX AF3-4 model underwent a second round of SREFLEX modelling to further evaluate the position/conformation of the first six nucleotides that appear as a 5′-overhang and not hybridized within the core RNA stem-loop. The SREFLEX AF3-4 model was divided into two rigid-bodies (5′-CCACAA-3′; 5′-GGUGCGGUGCUAACUAAAUCCCUAGCACCCC-3′) where the first 5′-CCACAA-3′ region underwent systematic manual rotations/translations to develop seven new SREFLEX AF3-4 based input templates, or alternatively these six 5′-nucleotides were swapped-out with the alternate AF3-predicted structures of the 5′-CCACAA-3′ region derived from AF models 0, 1, 2 and 3. SREFLEX was performed on the 11 new input RNA templates to develop representative individual RNA structures that fit the SAXS data.

CORAL rigid body modelling^104,105^ was used to generate models of the 2A protein and 2A-RNA complex. In both cases, the structural templates of the protein and RNA components were obtained from the X-ray crystal structure of the 2A-RNA complex (this study, PDB 9RVP). Unresolved RNA nucleotides (U31, A32 and A33) of the 2A-RNA complex were built using COOT^94^, by positioning a stereo-chemically ideal UAA trinucleotide between C30 and A34, linking the phosphodiester backbone manually and preforming real-space regularisation of the three modelled nucleotides using the 2Fo-Fc crystallographic map. The unresolved/likely flexible N- and C-terminal extensions of the 2A protein component were represented as dummy amino acids (N-terminus: GPLGSNPAS; C-terminus: HDVEMNPGEFPGRLERPHRD). CORAL was run five times against the respective 2A protein or 2A-RNA complex SAXS data to obtain a cohort of protein-alone or protein/RNA structures with alternative spatial positions of the 2A protein N- and C-terminal extensions. The final fit to the SAXS data for all atomistic models was performed using CRYSOL v2.8.3^115^ (30 harmonics, 256 points, constant enabled, average solvent electron density = 0.3416 *e*/Å^3^) and assessed by the reduced χ^2^ test or CorMap p-value. Additional details are provided in **Table 2**. All subtracted and un-subtracted SEC-SAXS 1D data frames, R_g_ correlations through the respective SEC-SAXS peaks, as well as MONSA, DAMMIN, SREFLEX and CORAL models and the corresponding model fits are made available in the Small Angle Scattering Biological Databank (SASBDB)^115^ under the accession codes: SASDWQ5 (2A protein), SASDWR5 (RNA) and SASDWS5 (2A-RNA complex).

### smFRET data acquisition and analysis

TMEV ‘end reporter’ (5ʹ Cy5, 3ʹ Cy3) and ‘internal reporter’ (5ʹ Cy5, U31 Cy3; oligos 2 and 3 detailed in **Table S2**) RNA oligonucleotides (IDT) were resuspended to 100 µM in nuclease-free water and stored at −80 °C prior to use. TMEV mutant ‘end reporter’ RNAs (oligos 16-19, detailed in **Table S2**) and EMCV ‘end reporter’ RNAs (oligos 20 and 21 detailed in **Table S2**) were prepared as described above. Samples were diluted to ∼12.5 pM RNA in 10 mM Tris-HCl pH 7.4, 150 mM NaCl, 1.0 mM DTT, 1.0 mM MgCl_2_ with 0.1 mg/mL BSA. For measurements with 2A protein, 0.5 µM of TMEV or EMCV 2A protein was added to the buffer above prior to RNA dilution as appropriate. smFRET data on freely diffusing molecules was acquired using a confocal-based single-molecule FRET spectrometer (EI-FLEX, Exciting Instruments) equipped with 520 and 638 nm laser lines operating under alternating laser excitation (ALEX). The ALEX cycle period was 100 µs (45 µs donor excitation, 5 µs lasers off, 45 µs acceptor excitation, 5 µs lasers off). 50 µL of sample was placed in a sealed coverslip (#1) chamber and data recorded for 2 hr at ambient temperature.

Triplicate measurements from independent experiments were collected and data combined. Analysis was performed in Anaconda 3 with Jupyter Notebooks using FRETBursts^116^ (v 0.8.3) as described previously^117^. Briefly, apparent smFRET efficiencies, defined as I_A_/(I_A_+I_D_), where I_D_ and I_A_ are the donor and sensitised acceptor emission intensities under 520 nm excitation were corrected with all four correction parameters as described previously^118^. For our system correction parameters where: α = 0.08319, δ = 0.06962, γ = 1.12903, β = 1.07552. Molecular stoichiometries, S, were evaluated via (I_D_ + I_A_) / (I_D_ + I_A_ + I_A_*), where I_A_* is the acceptor emission intensity upon direct excitation. After background estimation and subtraction, bursts were identified using an all photon sliding window algorithm as previously described^116^ with a photon rate of 10 photons and a minimum photon rate six times greater than the background rate. A burst size threshold of 10 and 20 photons was applied for the donor and acceptor channels respectively. Multimer events were removed from the donor channel by applying an upper limit threshold of 100 photons. Generally, 54% of molecules were donor only, 2% acceptor only and 44% doubly labelled. A dual channel burst search was used to extract doubly labelled bursts from each acquisition and find E and S with one or two Gaussian fits. The apparent donor-acceptor distance was calculated by 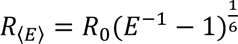 where *R*_0_ is 53 Å. We assumed random dipole orientation *κ² = ⅔* when approximating distances based on the peak FRET efficiency positions. Burst variance analysis (BVA) was performed as previously described^60^ and implemented in Jupyter Notebooks using FRETbursts^116^. Bursts were divided into sub-bursts (N = 7 photons) and the standard deviation calculated for each sub-burst population over the time period of individual bursts. Accessible volume (AV) modelling was performed using the FRET-restrained positioning and screening (FPS) tool^119^. The three radii model was used with dye parameters: R_1_ = 11 Å, R_2_ = 3 Å and R_3_ = 1.5 Å for Cy5 and R_1_ = 6.8 Å, R_2_ = 3 Å and R_3_ = 1.5 Å for Cy3. Both dyes were modelled with a linker length of 14 Å and linker width of 4.5 Å. Approximate distances from dye mean points were measured in PyMOL^97^.

### Electrophoretic Mobility Shift Assay (EMSA)

RNA oligonucleotides (oligos 4-15, detailed in **Table S2**) were reconstituted in nuclease-free water to 100 µM. RNAs were labelled at the 5ʹ end with Cy5-maleimide (GE Healthcare) or AzDye647-maleimide (Vector Labs) using the 5ʹ EndTag kit (Vector Labs) according to the manufacturer’s instructions, prior to phenol:chloroform extraction, ethanol precipitation and aqueous resuspension. The concentration of RNA was determined by measuring absorbance at 260 nm using a Nanodrop spectrophotometer (Thermo). 10 µL binding reactions contained 1.0 µL 500 nM fluorescently labelled RNA, 1.0 μL two-fold serially diluted 2A protein at concentrations of 0-12.5 µM in 10 mM Tris-HCl pH 7.4, 1.0 M NaCl, 1.0 mM DTT, 5.0 μL 2× buffer (20 mM Tris-HCl pH 7.4, 80 mM NaCl, 4.0 mM magnesium acetate, 2.0 mM DTT, 10% v/v glycerol and 0.02% w/v bromophenol blue) and 3.0 μl distilled water. Final concentrations in the binding reactions were 50 nM RNA, 1× buffer, a salt concentration of ∼140 mM and 2A protein concentrations of 0-1.25 μM. Samples were incubated at 37°C for 20 mins prior to analysis by non-denaturing 10% acrylamide/TBE PAGE (30 mins, 200 V constant). Gels were imaged with a Typhoon-5 using the 532 and 635 nm lasers/570BP20 and 670BP30 filters.

### Microscale Thermophoresis (MST)

Two-fold dilution series of TMEV 2A wild-type protein or mutants were prepared in SEC buffer (20 mM Tris-HCl pH 7.4, 1.0 M NaCl, 1.0 mM DTT) producing 2A concentrations ranging from 50 µM to 1.55 nM. For each binding experiment, fluorescently labelled RNA (oligos 4-15, detailed in **Table S2**) were diluted to 83.3 nm in MST buffer (20 mM Tris-HCl pH 7.4, 1.67 mM MgCl_2_, 1.0 mM DTT, 0.083% Tween-20) supplemented with 0.167 mg/ml bovine liver tRNA (Sigma). For the measurement, 2A ligand dilutions were mixed with labelled RNA in a 3:2 ratio, for a final concentration of 50 nM labelled RNA in 20 mM Tris-HCl pH 7.4, 400 mM NaCl, 1.0 mM MgCl_2_, 1.0 mM DTT, 0.05% Tween-20, 0.1 mg/ml tRNA and a 2A concentration series from 20 µM to 0.61 nM. The reaction was mixed by pipetting, incubated for at least 5 min at room temperature and then held at 4 °C. Capillary forces were used to load the samples into Monolith NT.115 Premium Capillaries (NanoTemper Technologies). Measurements were performed using a Monolith NT.115 instrument (NanoTemper Technologies) at an ambient temperature of 22 °C. Measurements were taken at 20-100% LED power and high MST power. An initial fluorescence scan was performed across the capillaries to determine the sample quality and afterwards 16 subsequent thermophoresis measurements were performed. Data analysis was carried out with an MST on-time of 1.5 seconds to determine binding affinities (MO.Affinity Analysis software, NanoTemper Technologies). Data were fitted to the K_D_ model (single-site binding) using MO.Affinity Analysis software (NanoTemper) using the following equation:

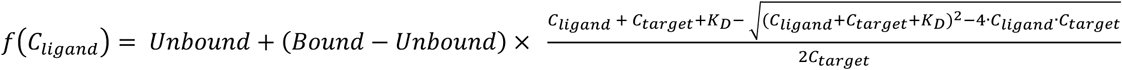

Where *f*(*C_ligand_*) is the normalised fluorescence value at a given ligand concentration *C_ligand_*. *Unbound* is the normalised fluorescence signal of the target alone, *Bound* is the normalised fluorescence signal of the complex. *K_D_* is the dissociation constant and *C_target_* is the final concentration of the target in the assay.

### In vitro translation assays

Plasmids were linearised with NdeI (NEB) prior to run off transcription with T7 RNA polymerase (prepared in-house). Messenger RNAs were recovered by phenol:chloroform extraction (1:1 v/v) and concentrated by ethanol precipitation before aqueous resuspension. Further purification was performed using the RNA clean and concentrator-5 kit (Zymo Research) as per manufacturer’s instructions. mRNA purity was assessed by gel electrophoresis and concentration determined by absorbance at 260 nm.

Messenger RNAs were translated in Flexi® rabbit reticulocyte lysate (RRL) (Promega). Typical reactions (10 µL) were composed of 66% (v/v) RRL, 25 µM amino acids (lacking methionine) and 0.2 MBq [^35^S]-methionine (Hartmann Analytic). The mixture was pre-equilibrated for 10 mins at 30 °C to allow for tRNA charging, prior to addition of the mRNAs and 2A protein to start the reaction. Translation reactions were conducted for 1 hr (30 °C), then stopped by addition of 100 µg/µL RNase A in 10 mM EDTA pH 8.0 (20 min, room temperature), followed by addition of 10 volumes of 2x Laemmli’s sample buffer, and heating (5 min, 95°C). Translation products were assessed by 15% SDS-PAGE. Dried gels were exposed to a Storage Phosphor Screen (PerkinElmer) and the screen scanned in a Typhoon-5 (Cytiva) using Phosphor imaging mode. Frameshift assays were performed in triplicate unless otherwise stated.

### Dual luciferase assays in cells

Adherent BHK-21 and HEK293T cells were transfected after seeding in 24-well plates to achieve 50% confluency. BHK-21 cells were transfected with 625 ng total plasmids at equimolar concentrations (250 ng pSGDlucV3.0 reporter and 325 ng pCEP4 expressing 2A) using TransIT-LT1 transfection reagent (Mirus Bio). HEK293T cells were transfected with 1 µg total plasmids in 1:1 ratio of pSGDlucV3.0 reporter to pCEP4 2A using PEI transfection reagent (MedChemExpress) as per manufactures instructions. BHK-21 cells were harvested 36 hpt, whereas HEK293T cells were harvested 24 hpt. Cells were washed with PBS and lysed with 1× passive lysis buffer (Promega), prior to luciferase activity measurement using either the Dual-glo® (HEK293T) or the Dual-luciferase® (BHK-21) reporter assay kits (Promega) according to the manufacturer’s instructions. Measurements were recorded using a GloMax Navigator microplate luminometer (Promega). The entire assay, from transfection to measurement, was performed in quadruplicate (BHK-21 cells) or triplicate (HEK293T cells).

## QUANTIFICATION AND STATISTICAL ANALYSIS

Crystallographic calculations (integration, scaling, merging) were performed as described in the methods text using the default software parameters unless otherwise stated. Processing and refinement statistics are detailed in **Table 1**.

SEC-SAXS calculations are described in methods text and detailed in **Table 2**. Additional information can be found in the Small Angle Scattering Biological Data Bank (SASBDB) under accession codes SASDWQ5 (2A protein only), SASDWR5 (RNA only) and SASDWS5 (2A-RNA complex).

smFRET data analysis was performed as described in methods text. Example Jupyter Notebooks for data analysis are deposited in Mendeley Data **10.17632/78kj389b6v.1**. FRET efficiencies are presented in the text as the mean ± SD. Data was collected and combined from three independent replicates. n indicates the number of bursts analysed after a dual channel burst search extracted doubly labelled molecules from each dataset.

For *in vitro* translation assays autoradiographs were quantified using ImageStudio 6.1 (LICORbio). The calculations of −1 frameshifting efficiency accounted for the difference in methionine content in the 0 and −1 frame products using the following equation:

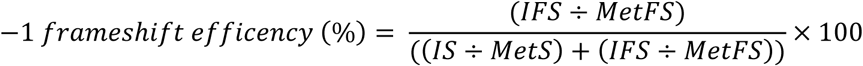

Where the number of methionines in the 0 frame and −1 frame products are denoted as MetS and MetFS respectively. The densitometry values for the same products are denoted by IS and IFS respectively. Frameshift assays performed in triplicate unless otherwise stated, where data is presented as the mean ± SD.

Frameshifting efficiencies in cells were determined by calculating the relative *Renilla*/firefly luciferase activities from each test construct and dividing by the relative luciferase activity from in-frame control (ifc) constructs, as described in the equation below:

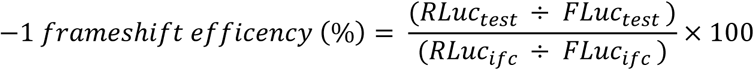

A one-way ANOVA followed by Tukey HDS test was used to test for significant differences in the mean −1 frameshifting efficiency of each test construct compared to wild-type RNA with wild-type 2A.

Graphing, data and statistical analysis was performed in Python using Spyder (v5.4.3). For statistical analysis the python package SciPy (v1.10.1) was used. Further statistical details can be found in figure legends and in the tables. For all figure legends, n is indicative of the number of independent experimental replicates conducted.

