## Supplementary Material for "A protein-dependent riboswitch activates ribosomal frameshifting in cardioviruses"

**Movie S1**. An annotated video highlighting key structural features of TMEV 2A protein bound to pseudoknot RNA. https://www.youtube.com/watch?v=I3jj-RdrPP8 (Related to **Figure 2**).

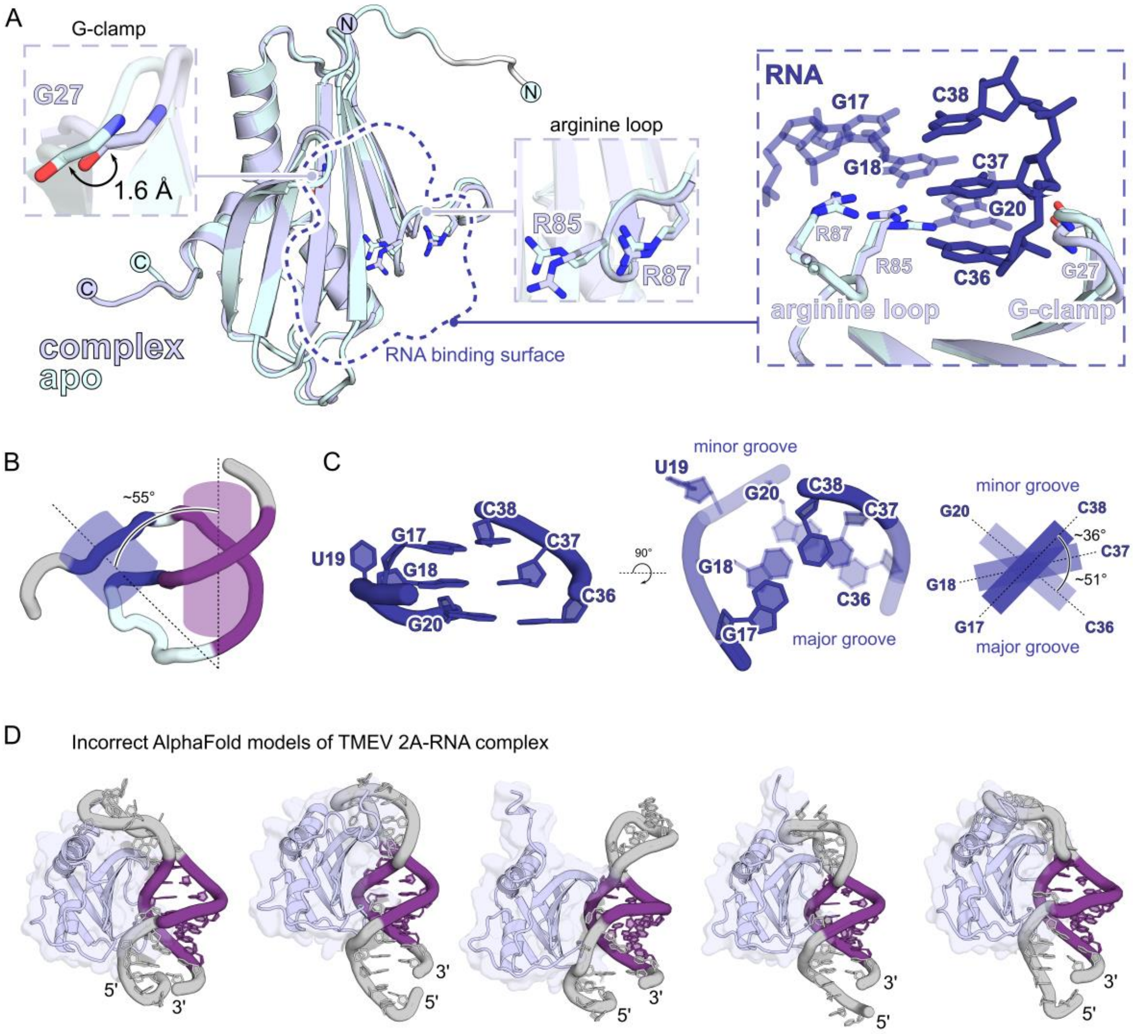


**Figure S1**. Comparison of the *apo* and *holo* TMEV 2A protein structures. Related to **Figures 1 and 2,** and **Table 1**.

A) Alignment of the *apo* (PDB 7NBV [1]) and *holo* (this study, PDB 9RVP) 2A protein structures reveals modest conformational plasticity of the 'G-clamp' (G27) and arginine loop (R85/R87) involved in RNA recognition. (*Insets*) Close-up of the 'G-clamp' and arginine loop shown as sticks.

B) Overview of the RNA pseudoknot showing the non-coaxial stacking of the two stems. The helical axes are represented by dotted black line and cartoon cylinders. The angle between the helical axis of S1 and S2 is ~55°.

C) Details of the non-consecutive guanine nucleotides in S1 that creates an unusual helical geometry. (*Left*) Side view showing the planarity of the three G-C base pairs. (*Middle*) Top view showing the effect of the flipped base U19 to the helical twist of S1. (*Right*) Schematic of S1 along the helical axes with helical twist angles indicated. The first and second base-pairs (G17-C38 and G18-C37) have a twist of ~36° whereas the twist of the second and third base-pair (G18-C37 and G20-C36) is ~51°. Helical twist angles analysed using the web 3DNA server [2].

D) AlphaFold3 [3] predictions of the 2A-RNA complex. An incorrect side-by-side arrangement is predicted in which 2A interacts with S2.

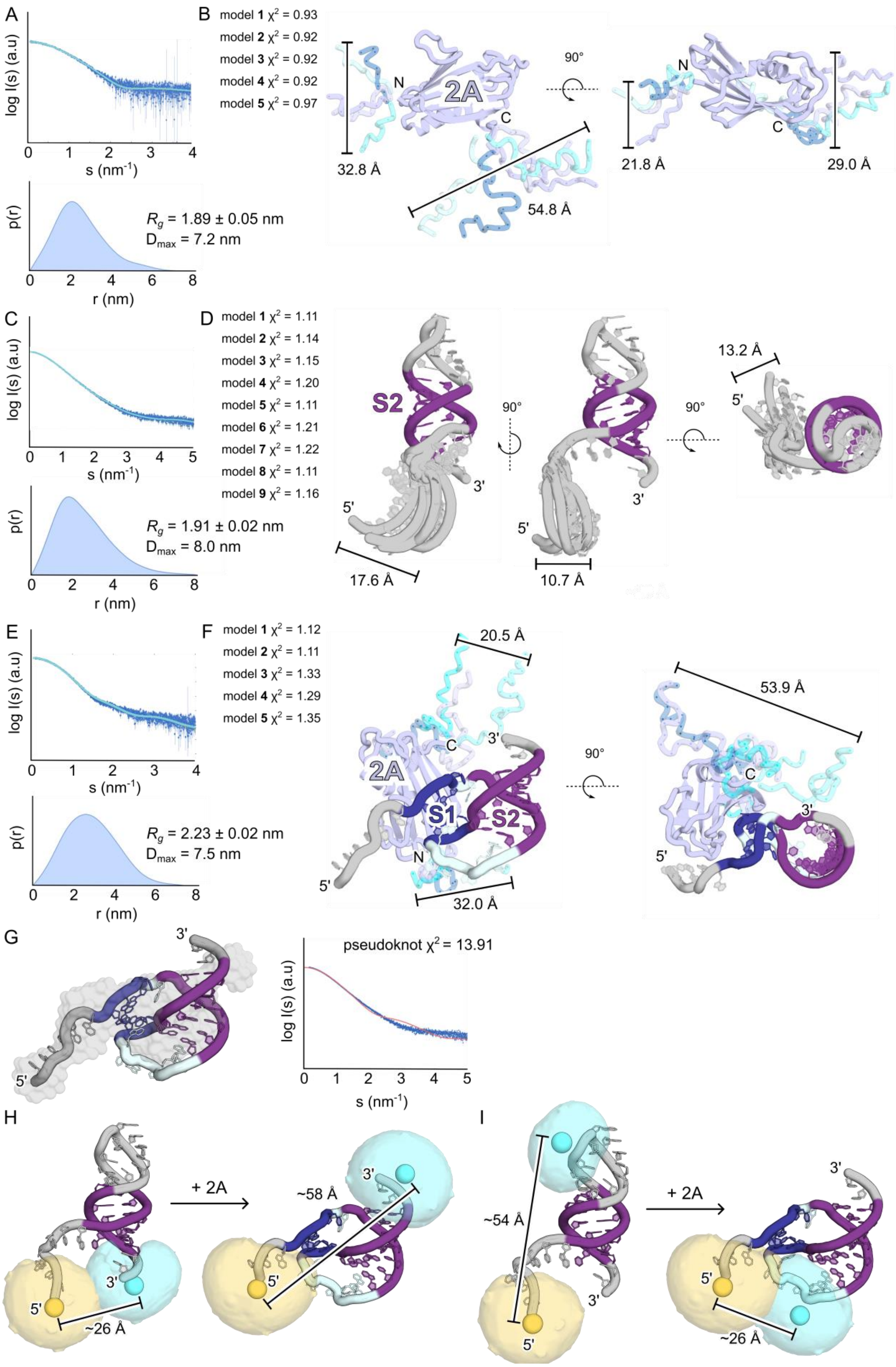
A
log I(s) (a.u)
s (nm-1)
p(r)
r (nm)
Rg = 1.89 ± 0.05 nm
Dmax = 7.2 nm
B
model 1 χ2 = 0.93
model 2 χ2 = 0.92
model 3 χ2 = 0.92
model 4 χ2 = 0.92
model 5 χ2 = 0.97
2A
N
C
32.8 Å
54.8 Å
90°
21.8 Å
29.0 Å
C
log I(s) (a.u)
s (nm-1)
p(r)
r (nm)
Rg = 1.91 ± 0.02 nm
Dmax = 8.0 nm
D
model 1 χ2 = 1.11
model 2 χ2 = 1.14
model 3 χ2 = 1.15
model 4 χ2 = 1.20
model 5 χ2 = 1.11
model 6 χ2 = 1.21
model 7 χ2 = 1.22
model 8 χ2 = 1.11
model 9 χ2 = 1.16
S2
5'
3'
17.6 Å
10.7 Å
13.2 Å
90°
E
log I(s) (a.u)
s (nm-1)
p(r)
r (nm)
Rg = 2.23 ± 0.02 nm
Dmax = 7.5 nm
F
model 1 χ2 = 1.12
model 2 χ2 = 1.11
model 3 χ2 = 1.33
model 4 χ2 = 1.29
model 5 χ2 = 1.35
20.5 Å
2A
S1
S2
N
C
32.0 Å
53.9 Å
90°
G
pseudoknot χ2 = 13.91
log I(s) (a.u)
s (nm-1)
H
+ 2A
~58 Å
~26 Å
I
~54 Å
+ 2A
~26 Å

**Figure S2 (previous page)**. Aligned atomistic models consistent with solution SAXS data for 2A protein, RNA, and 2A-RNA complex, demonstrating principal modes of flexibility. Related to **Figure 3** and **Table 2**.

A) Plots showing alignment between SAXS data for the 2A protein (dark blue circles) and calculated data from structural models (cyan lines), and the pair-distance distribution p(r) allowing the determination of $R_g$ and $D_{max}$. Mean $R_g$ values in nm are reported ± SD.

B) Flexible N- and C-termini of 2A (shades of blue) were modelled in CORAL [4]. Five superposed atomistic models are presented that fit the SAXS data, as assessed by reduced $\chi^2$ test. Displacements between equivalent termini are indicated in angstroms (Å).

C) Plots as in A) for RNA alone. Mean $R_g$ values in nm are reported ± SD.

D) Normal mode analysis using SREFLEX [5] illustrates conformational flexibility of the 5′ extension. Nine superposed atomistic models are presented that fit the SAXS data, as assessed by reduced $\chi^2$ test. Displacements between equivalent 5′ nucleotides are indicated in angstroms (Å).

E) Plots as in A) for 2A-RNA complex. Mean $R_g$ values in nm are reported ± SD.

F) Flexible N- and C-termini (shades of blue) of 2A were modelled in CORAL [4]. Five superposed atomistic models are presented that fit the SAXS data, as assessed by reduced $\chi^2$ test. Displacements between equivalent termini are indicated in angstroms (Å).

G) The pseudoknot conformation observed in the crystal structure does not explain the RNA-only scattering data, as assessed by reduced $\chi^2$ test. (*Left*) Pseudoknot model from crystal structure docked into SEC-SAXS envelope from RNA-only scattering data. (*Right*) Plots showing alignment between SAXS data for the RNA alone (dark blue circles) and calculated data from pseudoknot model (red line).

H, I) Accessible volume (AV) simulations calculated using FRET-restrained positioning and screening (FPS) tool [6] of RNA reporter constructs used in smFRET experiments. AV for Cy5 (yellow) and Cy3 (cyan) are represented as transparent surface and mean dye points as spheres. Distances from mean dye points were measured in PyMol [7]. H) end reporter. I) internal reporter.

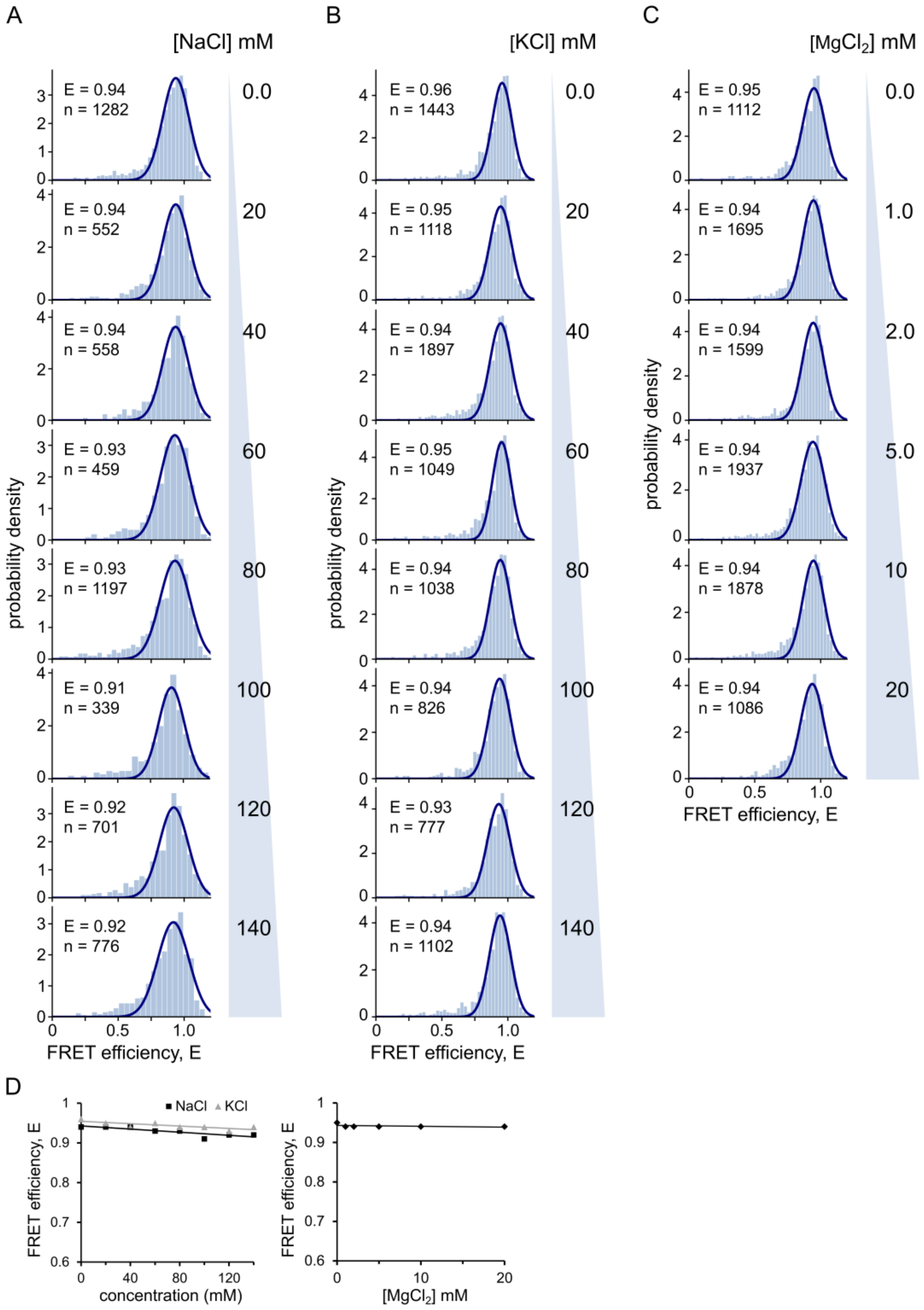


**Figure S3**. Ionic strength and identity have no effect on the conformational of the RNA. Related to **Figure 3**.

A-C) FRET efficiency (E) histograms of the 'end reporter' TMEV RNA fitted to a single Gaussian distribution. Corrected FRET efficiencies (E) and number of molecules (n) reported in each histogram. Titrations of A) NaCl B) KCl C) $MgCl_2$.

D) Summary of the 2D histograms with the corrected FRET efficiency plotted against ionic concentration. NaCl is represented by black squares and KCl by grey triangles.

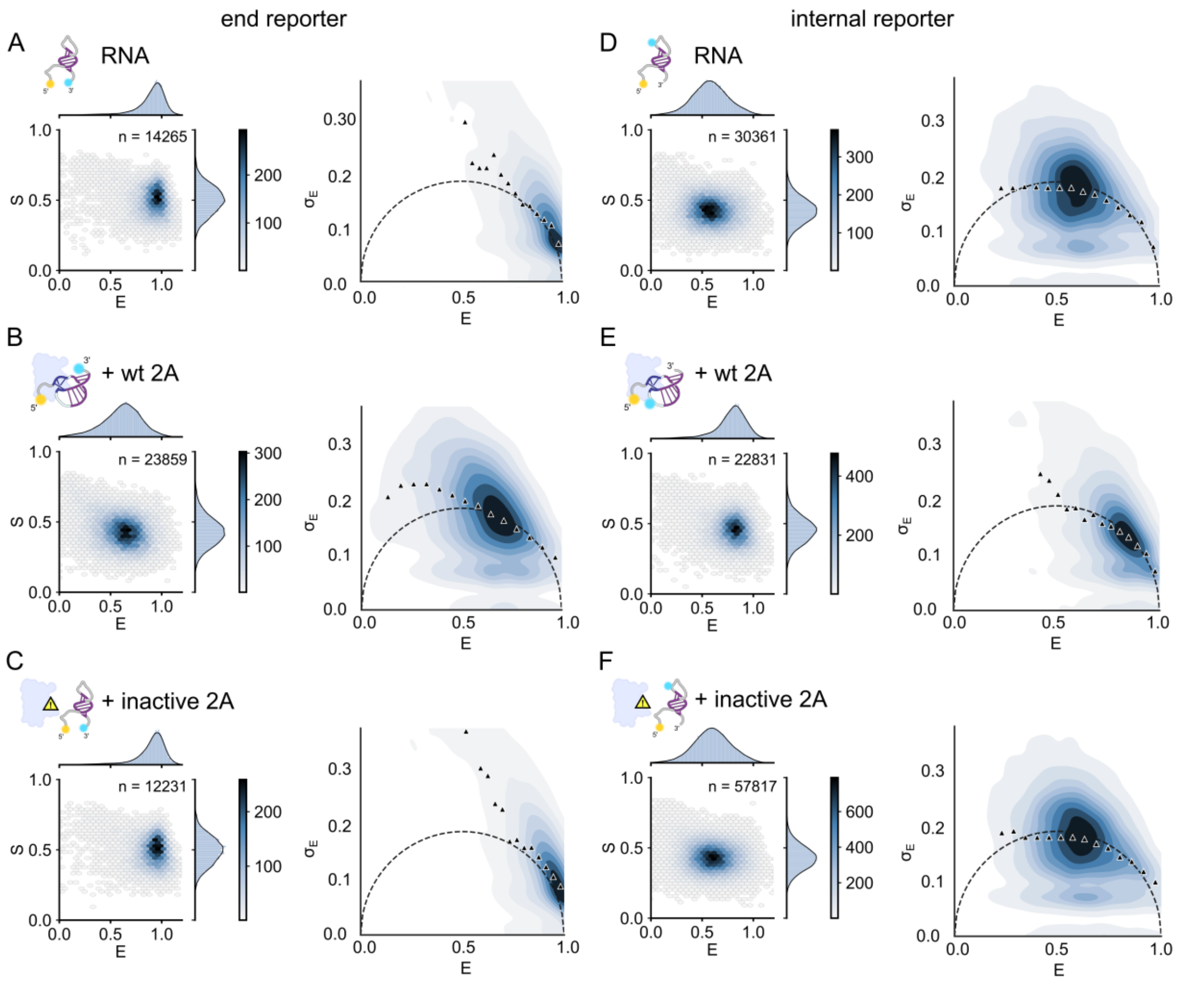


**Figure S4**. The RNA does not spontaneously convert between multiple stable conformations. Related to **Figure 3**.

(*Left*) 2D ALEX histograms of stoichiometry (S) against FRET efficiency (E) after filtering out the donor and acceptor only populations. The number of molecules in each dataset (n) is indicated. Data was collected from three independent experiments and combined. (*Right*) Corresponding burst variance analysis (BVA) contour plots assessing conformational dynamics of the RNA stimulatory element. Bursts were divided into sub-bursts of seven photons and the standard deviation calculated for each sub-burst population, indicated by black triangles. A standard deviation of sub-bursts greater than the theoretical shot-noise-limited variance (dashed black curve) suggests dynamics.

A-C) end reporter RNA (5' Cy5 and 3' Cy3). D-F) internal reporter RNA (5' Cy5 and Cy3 positioned on U31). A,D) RNA reporters in isolation. B,E) in the presence of 0.5 μM wild type 2A. C,F) in the presence of 0.5 μM inactive $2A_{R85A/R87A}$.

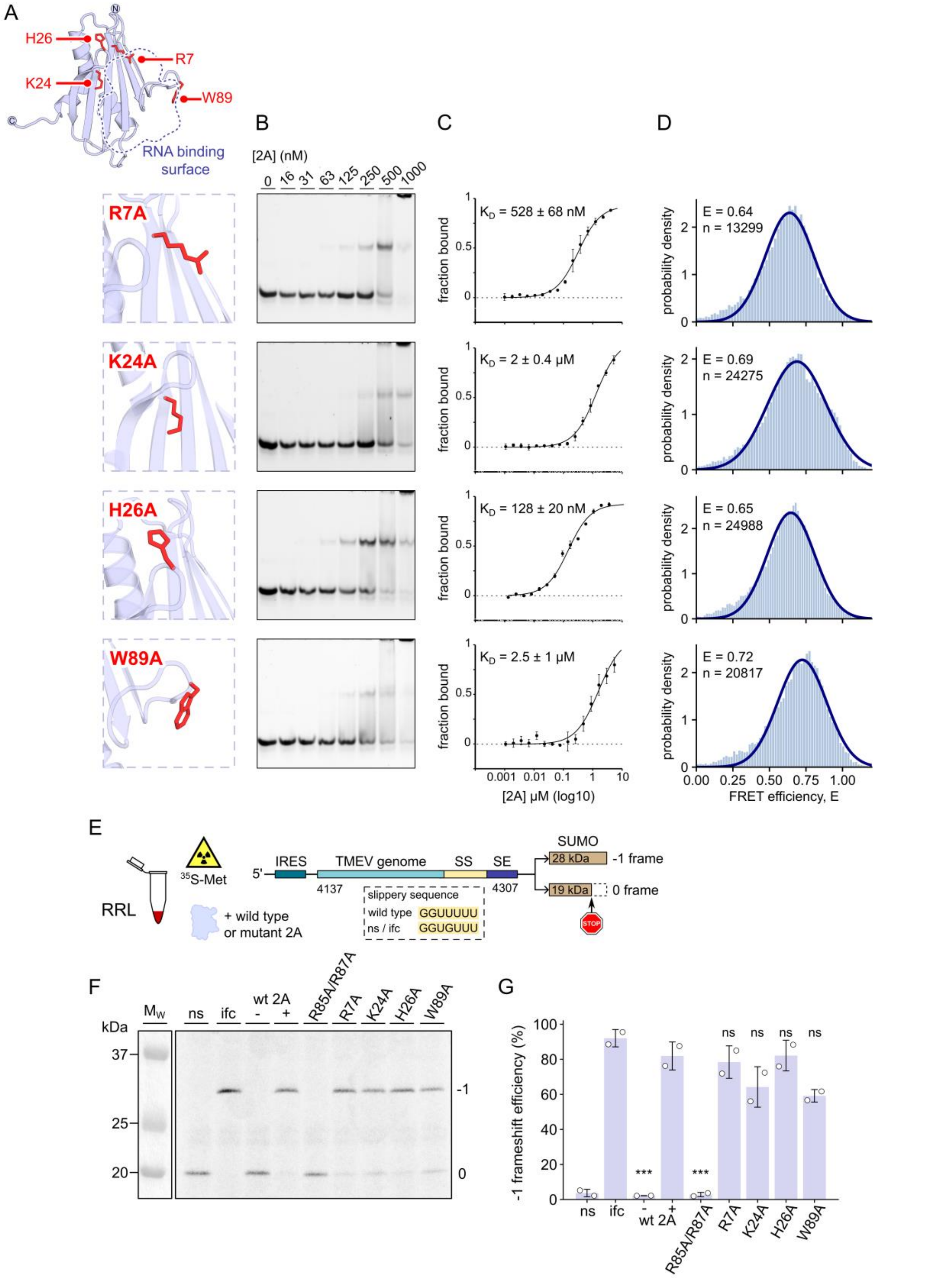


**Figure S5**. Mutagenesis of protein residues at the RNA binding interface has little effect on RNA recognition. Related to **Figures 4 and 5**.

A) Mutagenesis of residues in TMEV 2A that are involved in RNA binding. The locations of the mutations R7A, K24A, H26A and W89A are shown as red sticks. The RNA binding surface is represented as a blue dashed line.

B) EMSA analysis of the effects of mutant 2A proteins on RNA stimulatory element binding. Experiments were performed with 50 nM wild type RNA labelled with Cy5 at the 5' end. 2A concentrations were assessed between 0-1.0 μM. Following non-denaturing gel electrophoresis the gels were imaged with a Typhoon-5 scanner.

C) MST binding curves and approximate $K_D$ values using the same mutant 2A proteins. The $K_D$ fit model was used to describe a 1:1 binding stoichiometry. Error bars indicate standard deviation from the mean of three biological repeats.

D) Combined FRET efficiency (E) histograms of end reporter wild type RNA with mutant 2A proteins from three biological repeats. The distribution is fitted to a Gaussian distribution where $R^2 = 0.98$. Average FRET efficiency (E) and number of molecules (n) is reported on each histogram.

E) Schematic diagram of *in vitro* translation assay used to assess the effects of 2A mutations on frameshifting efficiency. Reporter constructs contained nucleotides 4137-4307 of the TMEV genome encompassing 35 codons of the viral genome prior to the viral frameshift site (coloured in blue). The slippery sequence (SS, yellow) and stimulatory element (SE, dark blue) are highlighted. SUMO protein (brown) was used as the product of translation in either the 0 or -1 frames. If elongation continues in the 0 frame the resulting product is ~19 kDa. Shifting into the -1 frame results in a ~28 kDa product. The non-slip (ns) control mRNA contains a mutated slippery sequence to prevent shifting into the -1 frame. The in-frame control (ifc) mRNA introduces another nucleotide after the stimulatory element, to simulate the effect of 100% -1 frameshifting. Translation reactions were carried out in rabbit reticulocyte lysate (RRL) in the presence of $^{35}$S-Met, analysed by SDS-PAGE and visualised by autoradiography.

F) SDS-PAGE and autoradiography analysis of 2A mutations on frameshifting efficiency *in vitro.*

G) Densitometric quantification of the -1 frameshift efficiencies, corrected for methionine content. Data are represented as the mean ± SD (n = 2). A one-way ANOVA with Tukey HSD test was used to compare 2A mutants to wild type RNA + 2A. ns, not significant, ***$p \leq 0.001$.

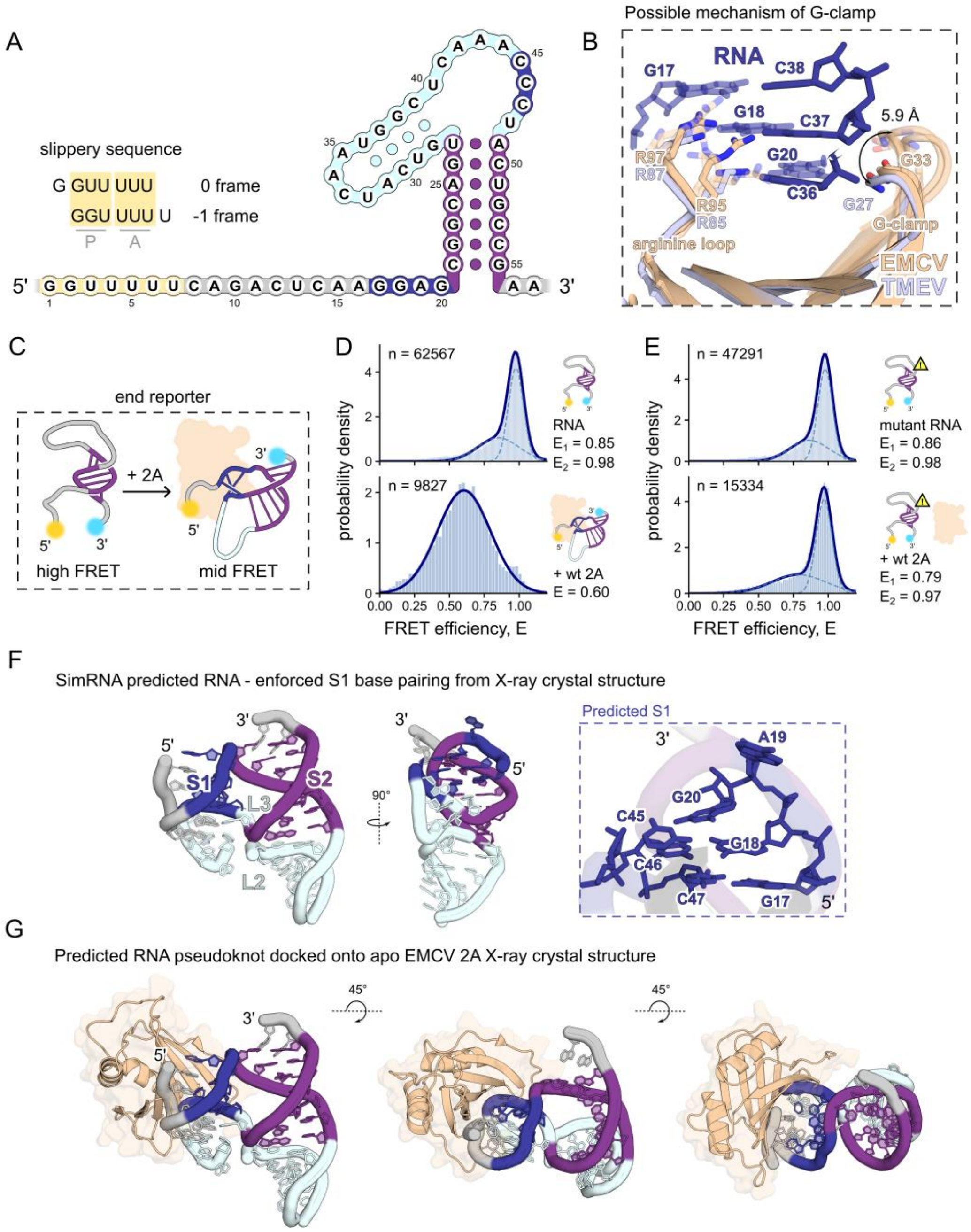


**Figure S6**. Proposed riboswitch mechanism in EMCV. Related to **Figures 1, 2 and 3.**

A) Schematic of the EMCV (47 nt) minimal RNA stimulatory element represented as a stem-loop. Slippery sequence is coloured in yellow, nucleotides not involved in base pairing are coloured in grey. Highly conserved GGxG and CCC motifs that comprise S1 coloured in dark blue, S2 coloured in purple and loops in cyan.

B) Alignment of *apo* EMCV crystal structure (wheat) (PDB 7BNY [8]) with TMEV 2A-RNA pseudoknot structure – this study (2A, light blue; RNA S1, dark blue). The RNA binding surface is likely to be conserved across the cardioviruses. NCS-related chains in the asymmetric unit of the *apo* EMCV 2A crystal structure (PDB 7BNY [8]) reveals a possible mechanism for the 'G-clamp' which demonstrates a pincer motion.

C) Schematic diagrams showing smFRET EMCV RNA end reporter construct design using the Cy3 (cyan) and Cy5 (yellow) FRET pair.

D) Corrected FRET efficiency (E) histograms, data combined from three biological repeats. Corrected average FRET efficiencies (E) and number of molecules (n) are reported in each histogram. Gaussian fits to our data are shown as dashed curves for two Gaussians and a solid blue line for a single Gaussian fit. $R^2$ = 0.99 for all Gaussian fits. Showing data for (*upper*) RNA in isolation, (*lower*) RNA in the presence of 0.5 µM wild type EMCV 2A.

E) As in D), but using EMCV C47G mutant RNA, predicted to prevent the formation of S1 (equivalent to TMEV C36G).

F) SimRNA [9] predicted model of the EMCV RNA pseudoknot with enforced S1 GGxG base pairing in two orthogonal views. (*Inset*) Close-up view of predicted S1 with flipped out base A19 in EMCV.

G) Predicted model of the EMCV 2A-RNA pseudoknot complex. Using the TMEV 2A-RNA pseudoknot complex as a reference, the *apo* EMCV 2A structure (PDB 7BNY) was aligned and docked onto the SimRNA [9] predicted EMCV RNA pseudoknot, suggesting a possible conserved mechanism of 2A-induced pseudoknot formation.

**Table S1**. DNA oligonucleotide sequences. Related to **STAR Methods**.

| Primer pair | Sequence | Source |
|---|---|---|
| **1** | Primers for subcloning TMEV 2A protein (wt and inactive 2A $_{R85A/R87A}$)<br>(F) 5' - AATTCATATGAATCCCGCTTCTCTCTACCG - 3'<br>(R) 3' - AATTGGATCCCTAGCCTGGGTTCATTTCTACATCATG - 5' | Integrated DNA Technologies (IDT) |
| **2** | Mutagenesis PCR primers – 2A$_{R7A}$<br>(F) 5' - CCCGCTTCTCTCTACGCCATTGATCTTTTCAT - 3'<br>(R) 3' - ATGAAAAGATCAATGGCGTAGAGAGAAGCGGG- 5' | Integrated DNA Technologies (IDT) |
| **3** | Mutagenesis PCR primers – 2A$_{K24A}$<br>(F) 5' - ATAACTTTTGACTACGCGGTCCACGGACGTCC- 3'<br>(R) 3' - GGACGTCCGTGGACCGCGTAGTCAAAAGTTAT- 5' | Integrated DNA Technologies (IDT) |
| **4** | Mutagenesis PCR primers – 2A$_{H26A}$<br>(F) 5' - TTTGACTACAAGGTCGCCGGACGTCCTGTGCT- 3'<br>(R) 3' - AGCACAGGACGTCCGGCGACCTTGTAGTCAAA- 5' | Integrated DNA Technologies (IDT) |
| **5** | Mutagenesis PCR primers – 2A$_{W89A}$<br>(F) 5' - GACGCTACCGCTCCGCGAAGAAGCCCATCCAC - 3'<br>(R) 3' - GTGGATGGGCTTCTTCGCGGAGCGGTAGCGTC - 5' | Integrated DNA Technologies (IDT) |
| **6** | Primers for cloning wild type and test sequences into pSGDlucV3.0<br>(F) 5' - CAGACTCGAGTCTCGAGTACACCACTTTCCG - 3'<br>(R) 3' - ACATAGATCTTTGTTTTGAATTCCTGCCTGG - 5' | Merck |
| **7** | Primers for cloning the in-frame control (ifc) into pSGDlucV3.0<br>(F) 5' - CAGACTCGAGTCTCGAGTACACCACTTTCCG - 3'<br>(R) 3' - ACATAGATCTGTTTTGAATTCCTGCCTGG - 5' | Merck |
| **8** | Primers for cloning - 1 bp stem 2 into pSGDlucV3.0<br>(F) 5' - CAGACTCGAGTCTCGAGTACACCACTTTCCG - 3'<br>(R) 3' - ACATAGATCTTTGTTTTTTGAATTCCTGCCTGG - 5' | Merck |
| **9** | Primers for cloning - 2 bp stem 2 into pSGDlucV3.0<br>(F) 5' - CAGACTCGAGTCTCGAGTACACCACTTTCCG - 3'<br>(R) 3' - ACATAGATCTTTGTTTTTTTTGAATTCCTGCCTGG - 5' | Merck |
| **10** | Primers for cloning TMEV 2A into pCEP4 for dual luciferase assays<br>(F) 5' - CATATGCTCGAGCCGCCACCATGAATCCCGCTTCTCTCTAC - 3'<br>(R) 3' - TTAGATCTGGATCCCTAGCCTGGGTTCATTTCTAC - 5' | Merck |

**Table S2**. RNA oligonucleotide sequences. Related to **STAR Methods**.

| Oligo number | Sequence | Source |
|---|---|---|
| **1** | RNA used in crystallisation of 2A-RNA complex<br>5' - CAAGGUGCGGUGCUAACUAAAUCCCUAGCACCCC - 3' | Integrated DNA Technologies (IDT) |
| **2** | Cy3/Cy5 ‘end reporter’ RNA used in smFRET experiments<br>5' - Cy5_CAAGGUGCGGUGCUAACUAAAUCCCUAGCACCCC_Cy3 - 3' | Integrated DNA Technologies (IDT) |
| **3** | Cy3/Cy5 ‘internal reporter’ RNA used in smFRET experiments<br>5' - Cy5_CAAGGUGCGGUGCUAAC(U-Cy3)AAAUCCCUAGCACCCC - 3' | Integrated DNA Technologies (IDT) |
| **4** | Stem 1 swap used in EMSA and MST<br>5' - CAACCUCCGGUGCUAACUAAAUGGGUAGCACCCC - 3' | Merck |
| **5** | AU stem 1 used in EMSA and MST<br>5' - CAAAAUACGGUGCUAACUAAAUUUUUAGCACCCC - 3' | Merck |
| **6** | UA stem 1 used in EMSA and MST<br>5' - CAAUUUUCGGUGCUAACUAAAUAAAUAGCACCCC - 3' | Merck |
| **7** | ΔU19 stem 1 used in EMSA and MST<br>5' - CAAGG_GCGGUGCUAACUAAAUCCCUAGCACCCC - 3' | Merck |
| **8** | Stem 2 swap used in EMSA and MST<br>5' - CAAGGUGCCCACGAUACUAAAUCCCAUCGUGGCC - 3' | Integrated DNA Technologies (IDT) |
| **9** | + 1 bp stem 2 used in EMSA and MST<br>5' - CAAGGUGGGGUGCUUACUAAAUCCCAAGCACCCC- 3' | Integrated DNA Technologies (IDT) |
| **10** | -1 bp stem 2 used in EMSA and MST<br>5' - CAAGGUGCGGUGCUACUAAAUCCCAGCACCCC- 3' | Integrated DNA Technologies (IDT) |
| **11** | -2 bp stem 2 used in EMSA and MST<br>5' - CAAGGUGCGGUGCACUAAAUCCCGCACCCC- 3' | Integrated DNA Technologies (IDT) |
| **12** | GGG mutant used in EMSA and MST<br>5' - CAAGGUGCGGUGCUAACUAAAUGGGUAGCACCCC - 3' | Merck |
| **13** | C38G mutant RNA used in EMSA and MST<br>5' - CAAGGUGCGGUGCUAACUAAAUCCGUAGCACCCC - 3' | Merck |
| **14** | C37G mutant RNA used in EMSA and MST<br>5' - CAAGGUGCGGUGCUAACUAAAUCGCUAGCACCCC - 3' | Merck |
| **15** | C36G mutant RNA used in EMSA and MST<br>5' - CAAGGUGCGGUGCUAACUAAAUGCCUAGCACCCC - 3' | Merck |
| **16** | Cy3/Cy5 GGG mutant RNA used in smFRET experiments<br>5' - Cy5_CAAGGUGCGGUGCUAACUAAAUGGGUAGCACCCC_Cy3 - 3' | Integrated DNA Technologies (IDT) |
| **17** | Cy3/Cy5 C38G mutant RNA used in smFRET experiments<br>5' - Cy5_CAAGGUGCGGUGCUAACUAAAUCCGUAGCACCCC _Cy3 - 3' | Integrated DNA Technologies (IDT) |
| **18** | Cy3/Cy5 C37G mutant RNA used in smFRET experiments<br>5' - Cy5_CAAGGUGCGGUGCUAACUAAAUCGCUAGCACCCC_Cy3 - 3' | Integrated DNA Technologies (IDT) |
| **19** | Cy3/Cy5 C36G mutant RNA used in smFRET experiments<br>5' - Cy5_CAAGGUGCGGUGCUAACUAAAUGCCUAGCACCCC_Cy3 - 3' | Integrated DNA Technologies (IDT) |

| | | |
|---|---|---|
| **20** | Cy3/Cy5 EMCV wt RNA used in smFRET experiments<br>5' -<br>Cy5_ACUCAAGGAGCGGCAGUGUCAUCAAUGGCUCAAACCC<br>UACUGCCGAA_Cy3 - 3' | Integrated DNA Technologies (IDT) |
| **21** | Cy3/Cy5 C47G mutant EMCV RNA used in smFRET experiments<br>5' -<br>Cy5_ACUCAAGGAGCGGCAGUGUCAUCAAUGGCUCAAACCG<br>UACUGCCGAA _Cy3 - 3' | Integrated DNA Technologies (IDT) |

**Table S3**. smFRET Gaussian fits parameters. Related to **Figure 3**.

| Sample | FRET $E_1$ | FWHM[a] $Gaussian_1$ | FRET $E_2$ | FWHM[a] $Gaussian_2$ |
|---|---|---|---|---|
| End reporter RNA | 0.85 ± 0.03 | 0.29 ± 0.03 | 0.96 ± 0.01 | 0.14 ± 0.01 |
| End reporter RNA + 2A | 0.63 ± 0.01 | 0.40 ± 0.01 | n/a | n/a |
| End reporter RNA + inactive 2A | 0.86 ± 0.02 | 0.31 ± 0.03 | 0.96 ± 0.00 | 0.15 ± 0.01 |
| Internal reporter RNA | 0.58 ± 0.00 | 0.40 ± 0.01 | n/a | n/a |
| Internal reporter RNA + 2A | 0.67 ± 0.02 | 0.47 ± 0.04 | 0.83 ± 0.01 | 0.23 ± 0.01 |
| Internal reporter RNA + inactive 2A | 0.60 ± 0.01 | 0.41 ± 0.01 | n/a | n/a |

[a]FWHM: full width half maxima. Data is presented as the mean ± standard deviation of three independent replicates.
